# Target Preference Maps: A machine learning model generalizing transferable drug-receptor interactions and guiding drug discovery

**DOI:** 10.1101/2025.08.01.668090

**Authors:** Filipe Menezes, Adam Wahida, Tony Fröhlich, Phillip Grass, Jan Zaucha, Valeria Napolitano, Till Siebenmorgen, Katarzyna Pustelny, Agata Barzowska-Gogola, Sarah Rioton, Kieran Didi, Michael Bronstein, Anna Czarna, Andreas Hochhaus, Oliver Plettenburg, Michael Sattler, Johannes Nissen-Meyer, Marcus Conrad, Razelle Kurzrock, Grzegorz M. Popowicz

## Abstract

Modern AI models can decode the genomic landscape and protein structure world. Yet, they fail to generalize to one of the most important fields: small-molecule drug discovery. Since the late 1970s, the advent of macromolecular crystallography inspired the notion that structural knowledge alone could enable a “lock-and-key” approach to drug design. However, drug discovery continues to depend on costly, resource-intensive, and largely serendipitous screening campaigns that probe only an infinitesimal fraction of the drug-like chemical space. Despite some successful cases, our understanding of, and reasoning from, non-bonded interaction chemistry remains limited for general applicability. Furthermore, though structural databases contain hundreds of thousands of entries, a strong historical bias pervades protein-drug structures, hindering reliable advances through AI scaling. Here, we present a machine-learning framework that learns atom-type-specific spatial preference maps from local protein microenvironments in protein-ligand structures. By excluding ligand topology from the model input and learning from local atom-level environments, the framework is designed to reduce dependence on whole-ligand memorization and to capture transferable interaction preferences. The resulting maps recover chemically meaningful interaction patterns, including cases involving bridging waters and metal-dependent environments. The model was validated using retrospective and prospective real-world data in drug optimization when targeting a challenging protein-protein interface. This shows that the method can provide interpretable workflows to guide molecule optimization and provide input for downstream generative or docking workflows.

## Introduction

Structure-based methods are central to modern drug discovery, supporting hit finding, pose analysis, and lead optimization. Physics-based and pharmacophore approaches, and, more recently, machine-learning methods, each capture useful aspects of protein-ligand recognition, but reliably transferring interaction patterns across diverse targets and chemotypes remains challenging. In most conventional structure-based workflows, drug-target interactions are approximated using simplified two-body functions, often derived from Newtonian physics or empirical measurements in model systems. The drug placement might also be proposed by finding a (local) minimum of these functions from random starting orientations. The induced-fit alterations, dynamics, and the context-dependent multi-body nature of ligand binding often limit the predictive accuracy of these physical models. Current machine learning (ML) models address it implicitly, by relying on hidden, learned representations rather than explicit interaction rules and hoping that model scaling will address remaining issues. The scaling, while successful in LLM, genomic, and even protein structure prediction, fails in the more diverse small-molecule datasets. This is because experimentally characterized molecules cover only an infinitesimally small fraction of the possible combinatorial space. Predictive performance declines when models are applied to structurally diverse or previously unseen complexes.^1–3^ The challenge is compounded by the training chemical space composition: its constituent molecules were either previously synthesized or a robust synthetic route to obtain them was already predicted.^4,5^ As a result, even extensive screening libraries remain concentrated in historically explored, accessible chemical spaces, shaped by prior synthetic campaigns and methodological bias, where a limited set of reactions often guides molecular design.^6^ This ultimately limits generalization and promotes overfitting, reducing the likelihood of identifying novel chemical matter with functional relevance and desirable novelty from both chemical and intellectual property perspectives.^7,8^ Consequently, the potential of structure-guided drug discovery remains difficult to realize in practice, and unsolvable by model scale alone. Although computational approaches are highly valuable, they are typically used alongside experimental Design-Make-Test-Analyze (DMTA) workflows rather than as replacements for them.^9^ A key challenge remains: to better relate receptor structure to ligand binding mode and interaction preferences, in ways that support DMTA efficiency while enabling exploration beyond historically overrepresented regions of chemical space. This remains an active area of research.^10,11^

Here, we show that an ML model can successfully generalize drug-receptor interaction principles from structural data alone. Inference of the presented model shows the ability to generalize beyond simple two-body approximations like H-bonding, lipophilic, or electrostatic attractions used in traditional methods. To achieve this, we train the model exclusively on local three-dimensional spatial features, abstracting ligand and receptor chemical connectivity (bond topology). The trained model infers three-dimensional probability distributions from empirical receptor structure data. By abstracting away from chemical connectivity, the model is prompted to capture local, spatial interaction patterns at the level of nearby atom groups. This supports generalization across diverse binding environments and differs from current structure-based models that condition on whole-ligand representations or directly optimize for pose- or affinity-related tasks.^12–14^ We reasoned that departing from learned whole-ligand representations could improve robustness to overfitting and reduce dependence on ligand-level memorization.^15^ We show that the model captures chemically meaningful, non-bonded interaction patterns and can even recover binding features associated with cofactors and bridging water molecules. In contrast with most established structure-based approaches, the model outputs continuous three-dimensional atom-type preference (probability) fields rather than discrete interaction features (atoms or pharmacophores) or scalar affinity scores. In addition to generalization of non-bonded chemistry, we show how the model can correctly infer transition metal chemistry, and we demonstrate experimentally how it suggests drug optimization modifications and provides selectivity-guided high-throughput screening.

The inferred preference fields, called Target Preference Maps (TPMs), share visual and conceptual relations to hotspot maps, pharmacophore fields, and knowledge-based interaction potentials, as they represent spatial interaction preferences within binding sites. They differ, however, in their origin: TPMs are generated *de novo* without any physical or chemical conditioning of the model. They are also distinguished by their features: rather than estimating energies, scoring poses, or encoding a small number of abstract interaction features (pharmacophores), TPMs predict continuous, atom-hybridization-state-resolved occupancy preferences from local structural environments. For simplicity we refer to it as an atom type that is not identical to a chemical element. Accordingly, TPMs are best viewed as interpretable interaction-preference models for ideal ligand design and hypothesis generation. Since the target learning objective of TPMs is probability fields (given this local arrangement, what ligand atoms best satisfy the environment), they also distinguish themselves clearly from machine-learned interatomic potentials (MLIPs),^16–21^ as the latter are approximations of the molecular potential energy surface.

### A Machine Learning Model to Capture Drug-Target Interactions

We built an ML model to learn non-bonded interaction preferences between ligand atom types and local receptor environments (**Figure 1A**). The model is specifically designed to address known limitations of structure-based drug discovery methods. ML models have previously shown the ability to capture the behavior of physical systems from limited sets of system-state parameters.^22,23^ In other words, they learned internal representations that sufficed to describe observed physical phenomena.

**Figure 1:**
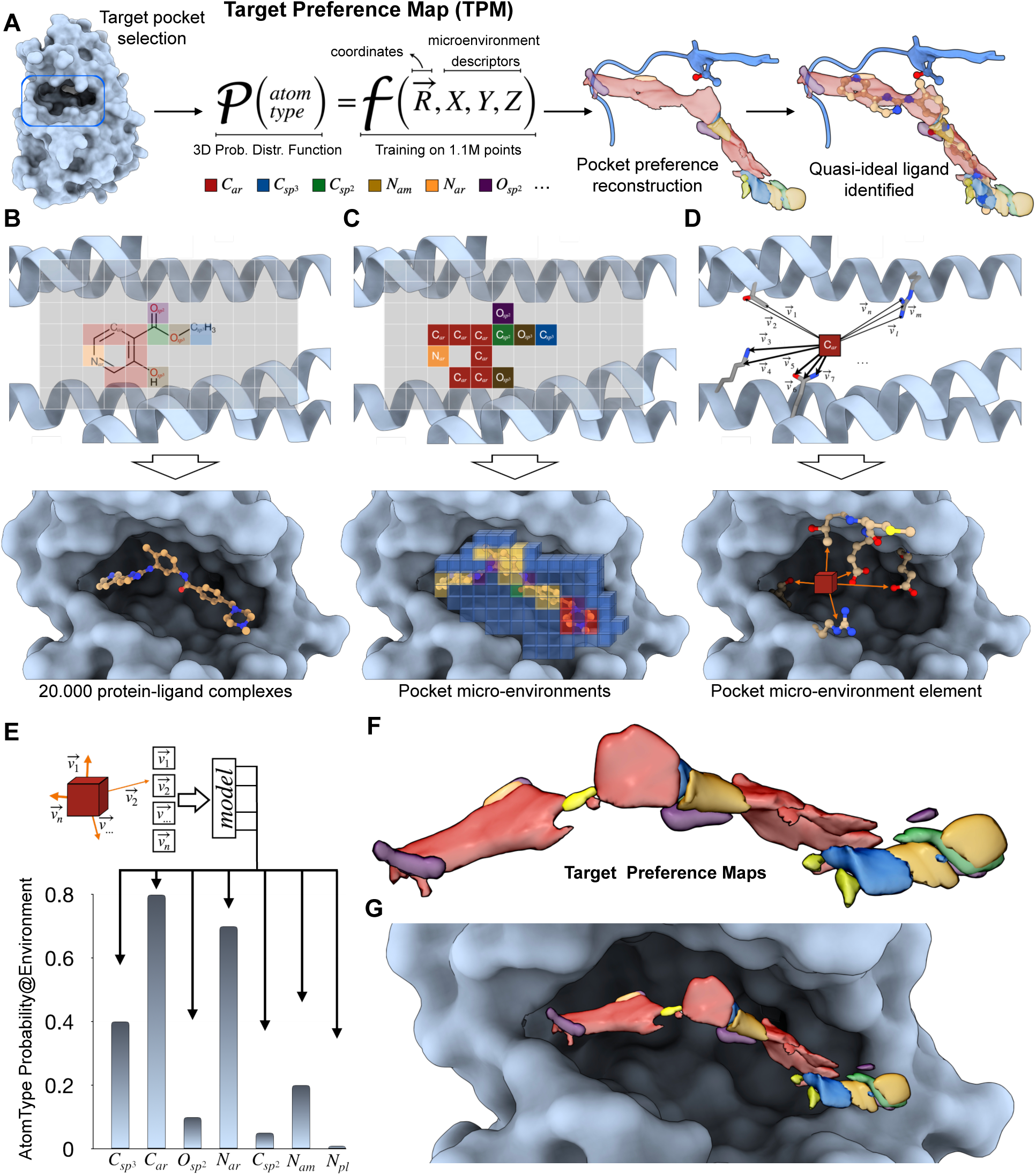
The Target Preference Map (TPM) Model. A) Conceptualization of the TPM model. Given the pocket of a biological target, we define atom-type-specific probability distribution functions. The function takes Cartesian coordinates and 3D chemical microenvironments (surrounding receptor atoms) as arguments and returns the likelihood of finding specific atom type in a receptor’s pocket. Reconstruction over the entire receptor pocket allows to identify quasi-ideal ligands. B-D) Two- (up) and three-dimensional (down) representation of the TPM-model construction. B) The biological target’s pocket is first voxelized, and then C) the atom types found in each element are identified. D) We then characterize the microenvironments by collecting all the nearby protein atoms in the form of neighbor vectors. E) The microenvironments are given as input to a transformer/perceiver model, which directly learns chemical matter distribution functions. F) Inference of the model assembled into three-dimensional clouds (TPMs), G) The same TPMs in the receptor pocket.

To allow this at a molecular level and minimize unwanted memorization, we hypothesized that the model should not directly access the drug’s chemical composition. This is because the landscape of drug-like molecules, estimated to encompass over 10^60^ entities,^24^ is sparsely explored by *ca.* 50,000 published protein-ligand complex structures.^6^ Instead, we trained the model to predict the atom type for each receptor volume element (**Figure 1B**). The atom type, composed of atom identity (*e.g.*, carbon), its hybridization state (*e.g.*, sp^3^; **Figure 1C**), and the surrounding receptor atoms together define a local microenvironment that constitutes one training point (**Figure 1D**). Each training instance, therefore, consists of a ligand atom type considered independently of the rest of the ligand, together with proximal protein atoms abstracted from the broader protein context. The experimentally observed occurrence of these local atom-type configurations is used as a binary indicator of their compatibility with productive binding contributions. This way, each drug-receptor complex structure yields 10^2^-10^3^ training points agnostic to ligand and receptor identity. The trained model can then be used to infer preferred atom types at points throughout the receptor binding site (**Figure 1E**), allowing the formulation of hypotheses about favorable local ligand chemistry.

Consequently, training becomes less dependent on drug-molecule connectivity, receptor identity, and ligand-specific imprinting in the pocket, although these sources of bias are reduced rather than fully eliminated. Further, abstracting chemical connectivity and exploring predictions for volumes smaller than an atom as microenvironments reduces limitations stemming from a shortage of experimental data: the 20,000 protein-ligand complexes^25,26^ used for training yield over 1,100,000 training points. A continuous assembly of such microenvironment predictions creates a volumetric map (TPM) that characterizes a receptor’s spatial preferences for different ligand atom types (**Figure 1F**). TPMs cover the entire investigated volume of the drug-receptor binding site (**Figure 1G**). Because each training point captures a local interaction preference, TPMs across atom types together describe a composite, ideal picture of the interaction features favored within the binding site. No real molecule can satisfy all such preferences simultaneously, because many preferred features are mutually exclusive; *e.g.*, aromatic nitrogens and sp^2^ oxygens can both support hydrogen bonding but cannot occupy the same position in a single chemical structure.

Nonetheless, TPM predictions reflect established non-bonded interaction patterns and recover binding features observed in real ligands. Because TPMs are trained on observed atom-type occupancies relative to background empty-space examples, ligands that satisfy more of these preferred local features may be expected to show improved binding compatibility, although TPMs are not intended as general affinity predictors. To balance the training dataset, we introduced datapoints corresponding to the absence of any ligand atom (NIL group). From the model perspective, these examples indicate positions where atom occupancy is less favored than empty space. **Figure 2** shows TPM predictions for several classes of biological targets. The use of local environments only allows the model to correctly infer binding features for every investigated structural class. Interestingly, TPMs can recover binding features associated with cofactors and bridging water molecules (see **Figure 4** and the respective discussion).

**Figure 2.**
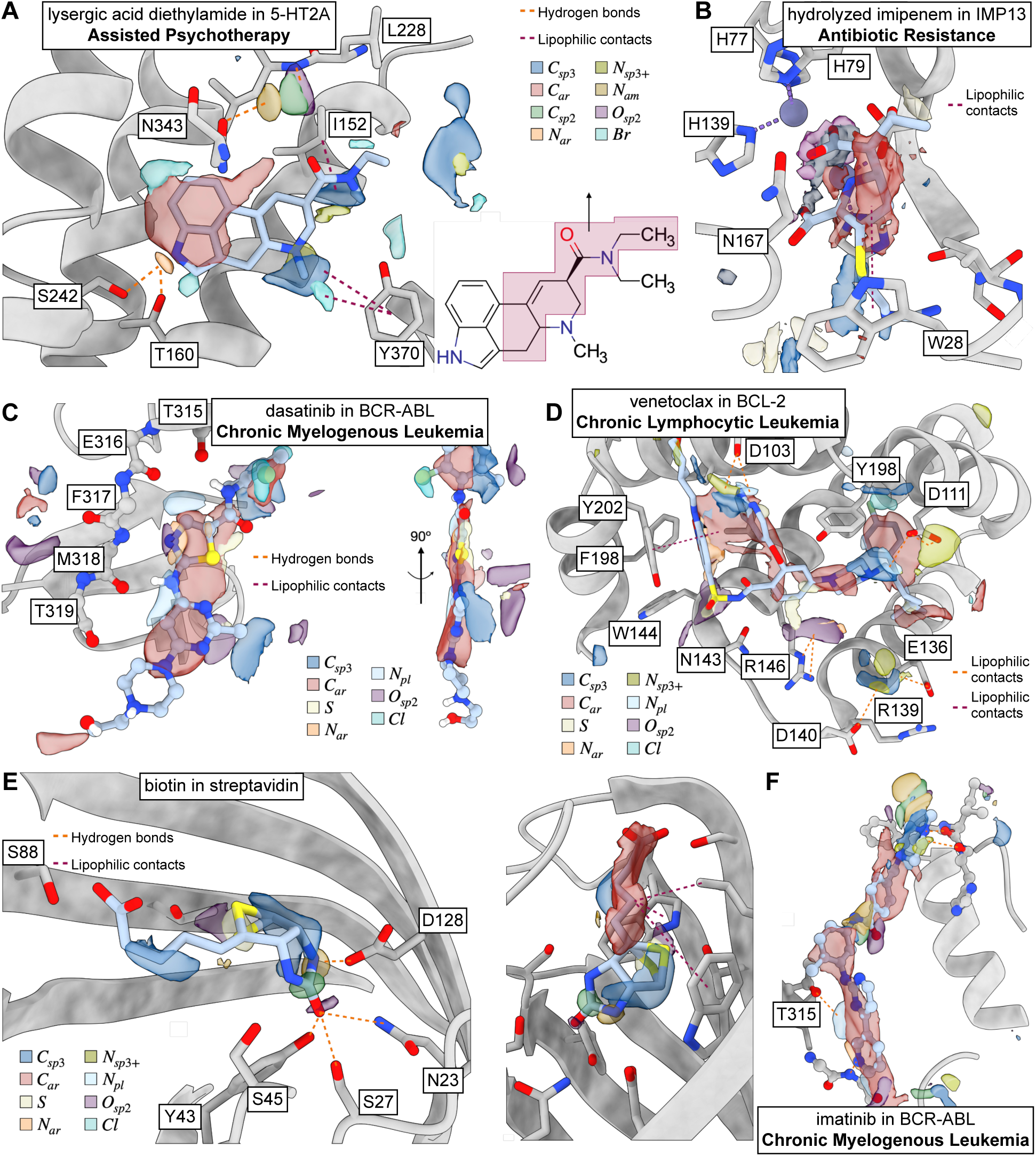
Examples of TPM Inferring Drug Features across Diverse Receptor Classes. A) Lysergic acid diethyl amide in the 5-HT2A G-Protein Coupled Receptor (GPCR), a model for assisted psychotherapy. TPM clouds predicted for the structure of the 5-HT2A GPCR with serotonin inside (PDB-ID 9ARX).^30^ Eight different clouds are represented in the structure. We observe an overlay of the aromatic clouds with the ligand’s aromatic groups. Similar observations apply to the aromatic nitrogen and the sp^3^ ammonium group. Lysergic acid’s amide group is uninvolved in any interaction with the protein, and this is reflected in the respective TPM predictions. Lastly, one of the bromine signals matches that of the bromolysergic acid diethyl amide ligand of this GPCR. B) Hydrolyzed imipenem in IMP13 (PDB-ID 6RZR).^31^ Although the model is oblivious to Zn atoms, it is likely inferred from side-chain arrangements around Zn atoms that a carboxyl group coordinates to the Zn^2+^ cations, as seen in known inhibitors. C) A type I kinase inhibitor, dasatinib, in its biological target BCR::ABL1 (PDB-ID 2GQG),^32^ a model for chronic myelogenous leukemia (CML). A good overlay of the aromatic clouds with the ligands’ aromatic groups and identification of hydrogen bond donors and acceptors to capture the kinase’s hinge region. Interestingly, the model overlays aromatic nitrogens and sp^2^ oxygens, despite their chemical incompatibility. This reflects the equivalence of these atom types in their hydrogen-bond accepting character. D) Venetoclax in BCL-2 (PDB-ID 6O0K),^33^ a model for chronic lymphocytic leukemia (CLL). The TPM model suggests adding ammonium groups close to some of the receptor’s asparagines (D103 and D111). Aromatic clouds are meant to capture π-stacking interactions with residues Y202 and Y198. E) Biotin in Streptavidin (PDB-ID 2F01).^34^ The model predicts adding aliphatic carbons, amide groups, and sulfur atoms in good agreement with the ligand’s structure. F) TPM predictions for imatinib in BCR::ABL1’s pocket (PDB-ID 2HYY),^35^ another model system for CML. TPMs capture the aromatic extent of the ligand, the amide group, and place aromatic and planar nitrogens to capture the kinase’s hinge by hydrogen bonding.

### The TPM Model Captures Non-Bonded Interaction Chemistry Patterns

We trained the TPM model on 30 atom types frequently observed in drug molecules. However, this framework is compatible with other atom-type partitionings, *e.g.*, based on quantum-chemical atomic properties. Our initial benchmark was designed to assess whether the model captures chemically meaningful non-bonded interaction patterns from the training data. We begin by evaluating the model’s accuracy (**Figure 3A**) and AUC (**Figure 3B**) relative to sample size for each atom type. Atom-type metrics appear largely independent of sample size (**Figures 3A, B**). Most atom types show good accuracy (>70%), with phosphorus exceeding 90% and boron nearing 95%. For boron, such high accuracy despite the small sample size (∼200 samples) may reflect the relatively constrained structural contexts in which boron atoms are observed in protein binding sites; *e.g.*, boronic-acid warheads are frequently used to target hydrolytic enzymes. For phosphorus, this likely reflects the abundance of phosphate groups in the training set (nucleotide sites). AUC values are also high overall (>75%), except for S_sp3_ and N_tr_. Again, boron and phosphorus scored the highest, with phosphorus showing a slightly better AUC. Atom types with lower accuracy despite large sample sizes prompted further examination. For example, O_sp2_ and O_sp3_ have similar sample sizes, yet O_sp2_ is predicted with ∼5% greater accuracy. Both atom types participate in polar, directional interactions, but O_sp2_ has typically lower partial charges (–0.5 electrons vs. –0.25 electrons), which may favor stronger contacts. Likewise, C_ar_ and N_ar_ show identical accuracy despite a 10-fold difference in sample size (200,000 vs. 20,000). N_ar_ commonly engages in more directional hydrogen bonding interactions, while C_ar_ often participates in less directional, lipophilic contacts and π-stacking, which may explain the model’s lower accuracy metrics. A final example contrasts C_ar_ and C_sp3_, which have similar sample sizes. Still, C_ar_ is predicted to be ∼5% more accurate, likely because C_sp3_ lacks π-stacking ability and predominantly participates in non-directional lipophilic interactions. These comparisons suggest that model performance is influenced more strongly by interaction type and directional nature than by sample count alone: atom types involved in polar and hydrogen bonding tend to be predicted more accurately. Conversely, performance is lower for atom types associated primarily with lipophilic interactions and scaffolding elements. This pattern differs from the behavior often emphasized in current ML folding frameworks, where shape and hydrophobic packing are more readily captured than polar and hydrogen-bond interactions.^27–29^ These results suggest that the available sample sizes were sufficient to reveal chemically meaningful differences across atom types, and that observed accuracy differences reflect interaction characteristics at least as much as training-set size.

**Figure 3.**
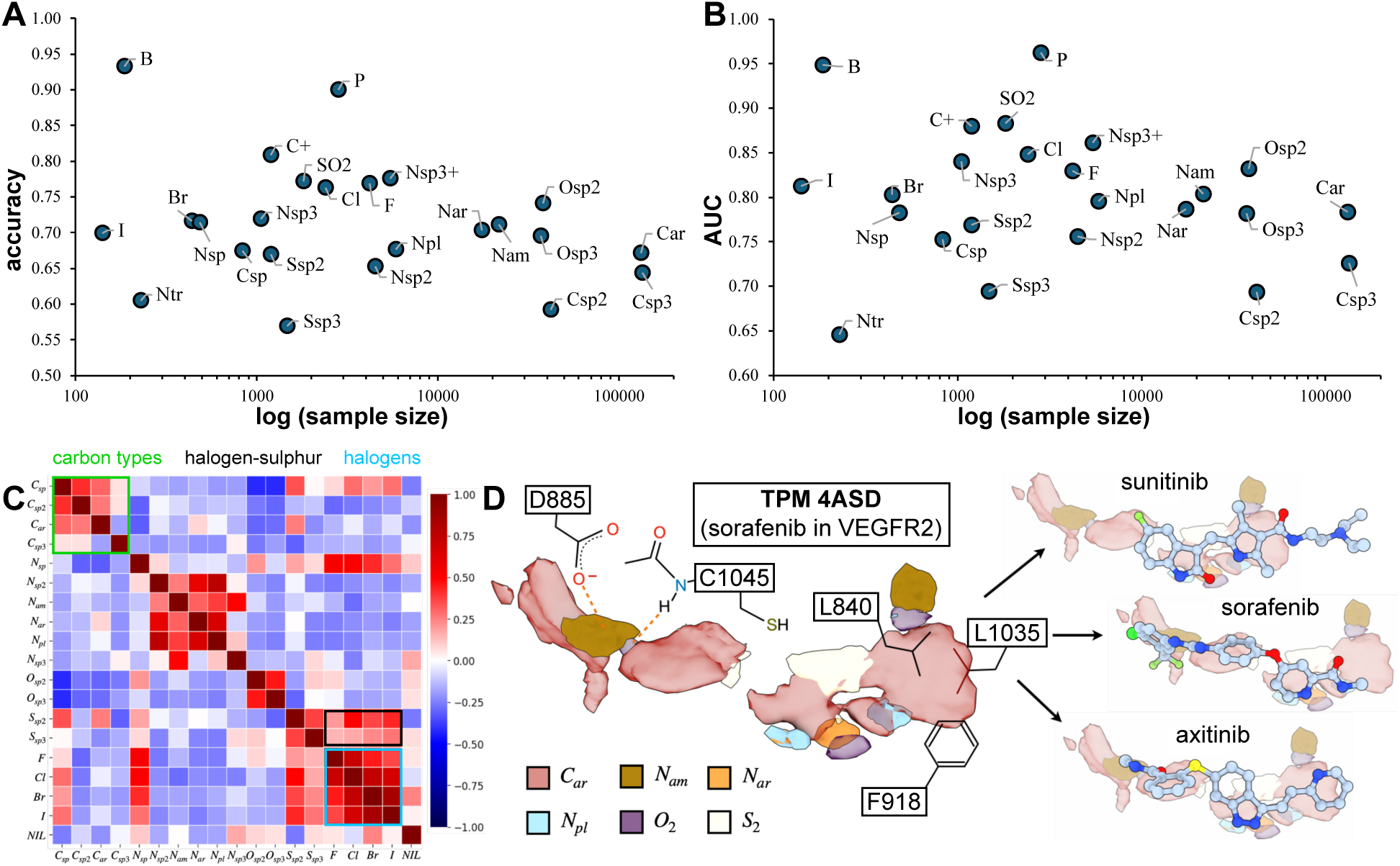
Statistics and Inference of the TPM Model. A. TPM-model accuracy for every atom type considered against the respective sample size on a logarithmic scale. B. TPM-model AUC for every atom type considered against the respective sample size on a logarithmic scale. These two blocks show that polar atom types may achieve higher accuracy than apolar ones given similar sample sizes, because non-bonded interactions are more directional. C) Correlation matrix of atom types: given that one atom type is found at a specific point in the target’s pocket, what is the likelihood of finding other, alternative atom types? Observed (anti)correlations align perfectly with traditional medicinal chemistry substitutions. D) Cross-validation of the TPM model on VEGFR with sorafenib (PDB-ID 4ASD), a model for lung cancer. Overlay of TPM predictions with the structure of other VEGFR inhibitors, sunitinib and axitinib. The TPMs show a good overlay with the chemical composition of axitinib, although the kinase’s pocket contained sorafenib. This indicates the model’s generalization capability. Noteworthy are the predictions to extend the aromatic density and the introduction of a sulfur atom, where axitinib contains a linker.

An additional evaluation concerns the extent to which atom types with similar properties are predicted at the same spatial locations. To examine this, we plotted the correlation between model-predicted atom types at each spatial point (**Figure 3C**). Atom types with related binding-relevant properties show strong correlations. For example, all unsaturated carbon types (C_sp2_, C_sp_, C_ar_) correlate with one another, but they do not correlate with saturated carbons (C_sp3_), devoid of π-electron systems. As expected, halogen atoms (F, Cl, Br, I) showed strong internal correlation. The small, electron-dense fluorine, the halogen atom least prone to forming dispersion (van der Waals) interactions, shows a weaker correlation with the later halogens (Br, I). Conversely, iodine shows a weaker correlation with earlier halogens (F, Cl), consistent with its greater lipophilicity. These changes in correlation are gradual, with chlorine and bromine showing intermediate behavior consistent with partial interchangeability with neighboring halogens. Among the halogens, bromine and iodine show the strongest apparent interchangeability, consistent with their higher lipophilicity. Halogens also correlate with sulfur atoms, again reflecting the high lipophilic character of these atom types. They also correlate well with N_sp_, as those found in nitrile moieties, consistent with the medicinal-chemistry intuition that nitriles can replace halogens in constrained pockets, most prominently, fluorine. A notable cross-correlation was observed between N_sp3_ and N_am_, consistent with their shared hydrogen-bonding roles. Interestingly, we also observe a correlation between O_sp2_ and N_sp_, likely reflecting their hydrogen-bond acceptor character. However, these two atom types are not fully interchangeable because C≡N and C=O groups differ in polarity, lipophilicity, and hydrogen-bond directionality. Oxygen predictions also show a weak correlation with the NIL maps, which may reflect positions compatible with water occupancy in the binding site.^36^

The observed anticorrelations are also informative: for example, oxygen atom (O_sp2_, O_sp3_) predictions are anticorrelated with many other atom types, particularly carbon. This is consistent with the fact that O_sp2_ and O_sp3_ commonly participate in hydrogen-bond networks, whereas carbon atoms generally do not. Together, these examples indicate that the TPM model captures distinctions beyond coarse pharmacophore categories such as donor/acceptor, aliphatic, and aromatic features, including differences associated with atomic hybridization and local electronic character. These observations support the conclusion that the model learns chemically meaningful local interaction preferences relevant to ligand binding.

### TPMs Recover Conserved Chemotypes Associated with Ligand Binding

We next examined to what extent the model captures transferable interaction preferences, rather than reproducing ligand-specific features imprinted by receptor conformational adaptation. To investigate this, we generated TPMs for one receptor using different co-crystals with different drugs. In all cases, the corresponding structural family was excluded from the training set. We used the structures of VEGFR2 with sorafenib, axitinib, and sunitinib as references (PDB IDs 4ASD, 4AGC, and 4AGD, respectively; **Figure 3D**).^37^ We began by analyzing the TPMs generated from the VEGFR model co-crystallized with sorafenib (4ASD). These TPMs contain features that overlap with structural elements present in the other two inhibitors, axitinib and sunitinib, among which, sunitinib shows the least overlap with the 4ASD TPMs. We initially considered whether this might reflect a distinction between type I kinase inhibitors, such as sunitinib and axitinib, and the type II inhibitor sorafenib. However, superposing the three PDB structures reveals substantial similarity in their VEGFR2 binding modes. This suggests that the remaining TPM differences reflect local differences in how each inhibitor engages the ATP-binding pocket.

Several structural elements are nonetheless conserved, and TPMs suggest how local receptor-drug interactions can be explored across chemotypes. For instance, an N_pl_ signal aligns with sunitinib’s pyrrole-containing oxoindole motif. The oxoindole oxygen also aligns with the O_sp2_ signal. Axitinib shows an even broader alignment with the 4ASD-TPMs. The amide signal (N_am_) overlaps with axitinib’s amide group, and the drug covers a larger extent of the C_ar_ densities. Importantly, the amide signal is also present in the TPMs calculated from sunitinib’s co-crystal structure, although sunitinib does not occupy that pocket subregion. The conservation of this signal across structures is consistent with the amide functionality representing a transferable local binding preference rather than a purely ligand-specific induced-fit effect. We also observe that axitinib’s indazole group aligns with N_pl_ and N_ar_ TPM features calculated from sorafenib’s co-crystal, even though sorafenib places a N_ar_ functionality in that region. Notably, no N_ar_ signal is detected near axitinib’s terminal pyridyl. As shown in **Figure S1A**, this group does not make a direct interaction with the protein in the VEGFR2 pocket. This observation is consistent with the view that the corresponding TPM features are inferred from local interaction patterns rather than directly copied from the bound ligand. Nevertheless, this ring extends into a region associated with aromatic preference in the 4ASD-derived TPMs. Finally, axitinib places a sulfur atom close to an S_sp3_ signal present in the sorafenib-derived TPMs, although sorafenib itself lacks a sulfur linker in that position. Figure S1 provides the corresponding analyses for the other two co-crystal structures.

Together, these analyses show that TPMs capture recurring local features associated with ligand binding. Because these features recur across structures containing different ligands, the results are consistent with reduced dependence on ligand-specific memorization and with learning of transferable local interaction preferences. This suggests that TPMs may help guide medicinal chemistry optimization toward new ligand hypotheses beyond those directly observed in the reference structure. TPM predictions span the investigated binding-site volume, including regions not occupied by the ligand present in the reference crystal structure. This can be useful for proposing extensions or substitutions during chemical optimization. As expected, some TPM features are mutually incompatible with the baseline ligand pose. This follows from the fact that no single molecule can simultaneously satisfy all preferred local features. For example, Figure S2 shows overlapping O_sp2_ and N_ar_ signals that correspond to alternative ways of satisfying a hinge-region hydrogen-bond interaction in a kinase pocket. Figure S2 further compares TPM signals for representative type I and type II kinase binders. TPMs generated from different type I inhibitors show substantial agreement. Expectedly, TPMs derived from type I and type II inhibitor complexes also differ in ways consistent with their distinct binding modes. We also generated TPMs for additional classes of drug targets. In these examples, the predicted maps show substantial agreement with features present in potent ligands (**Figures S3**, **S4**, and **S5**). In all cases, the models used for prediction were trained with the corresponding query structures excluded from the training set to reduce the risk of data leakage and direct memorization. Overall, these examples indicate that TPMs recover spatially organized chemotypes associated with experimentally observed ligand-binding modes. Additional benchmarks are presented in Figure S6 and its related content.

### TPM detects Transition Metal Interaction Patterns

We showed that the TPM model is sensitive to the chemical nature and spatial arrangement of receptor binding sites. Because of sparse training data, cofactors, including metals, were removed from the training set. However, cofactors can leave a prominent structural imprint on the surrounding receptor environment, and the model appears capable of capturing these aspects indirectly. To examine this, we investigated how TPM-density predictions behave in the presence of transition-metal-dependent binding environments, even though the model itself is agnostic to the metal atoms. We selected IMP-13, a metallo-β-lactamase involved in antibiotic resistance and a target for which no approved inhibitors are currently available.^31^ IMP-13 coordinates two adjacent zinc ions at its active site. These zinc atoms activate bound water molecules, generating a hydroxide nucleophile that hydrolyzes β-lactam antibiotics. This is illustrated in the crystal structure of apo-IMP-13 (PDB ID 6R78; **Figure 4A**). TPM predictions based on the structure with PDB ID 6R73, co-crystallized with hydrolyzed meropenem, show a prominent O^sp3^ signal at the location where a hydroxide anion bridges the two metal centers in related structures (**Figures 4B, C**). Notably, in 6R73 itself, no such bridging hydroxide is present, as the hydrolyzed antibiotic sterically and chemically disfavors hydroxide coordination. A prominent TPM density for carboxylate-like oxygen features is also observed at the binding site, overlapping closely with carboxylate groups found in other ligand-bound structures (6RZR, 6RZS, and 6S0H; **Figures 4D, E**). Because the reference co-crystal 6R73 does not contain zinc-bound carboxylates, this signal is more consistent with a transferable local structural preference than with direct copying of the bound ligand. The TPM model also suggests sulfur-containing features in proximity to the zinc ions (**Figure 4F**), in line with known aspects of zinc coordination chemistry. By contrast, no N_ar_ density is predicted directly around the metal centers (**Figure 4G**). One possible explanation is that the N_ar_/C_ar_ density is spatially displaced from the zinc sites, such that aromatic nitrogens would not provide an additional favorable interaction in that region. Another notable signal is the strong TPM density for ammonium-like groups (N_sp3_, **Figure 4H**; N_sp3+_, **Figure 4I**) extending toward the zinc ions. Although ammonium ions do not coordinate transition metals, their positive charge and local electrostatic character may partly explain this similarity in TPM signals. In our experience with the TPM model, this is one of the few cases in which N_sp3_, N_sp3+_, and sulfur-associated signals overlap so closely. To further assess the robustness of these observations, TPM densities were also calculated for an additional structural model, yielding consistent qualitative results (**Figures 4J-O**).

**Figure 4.**
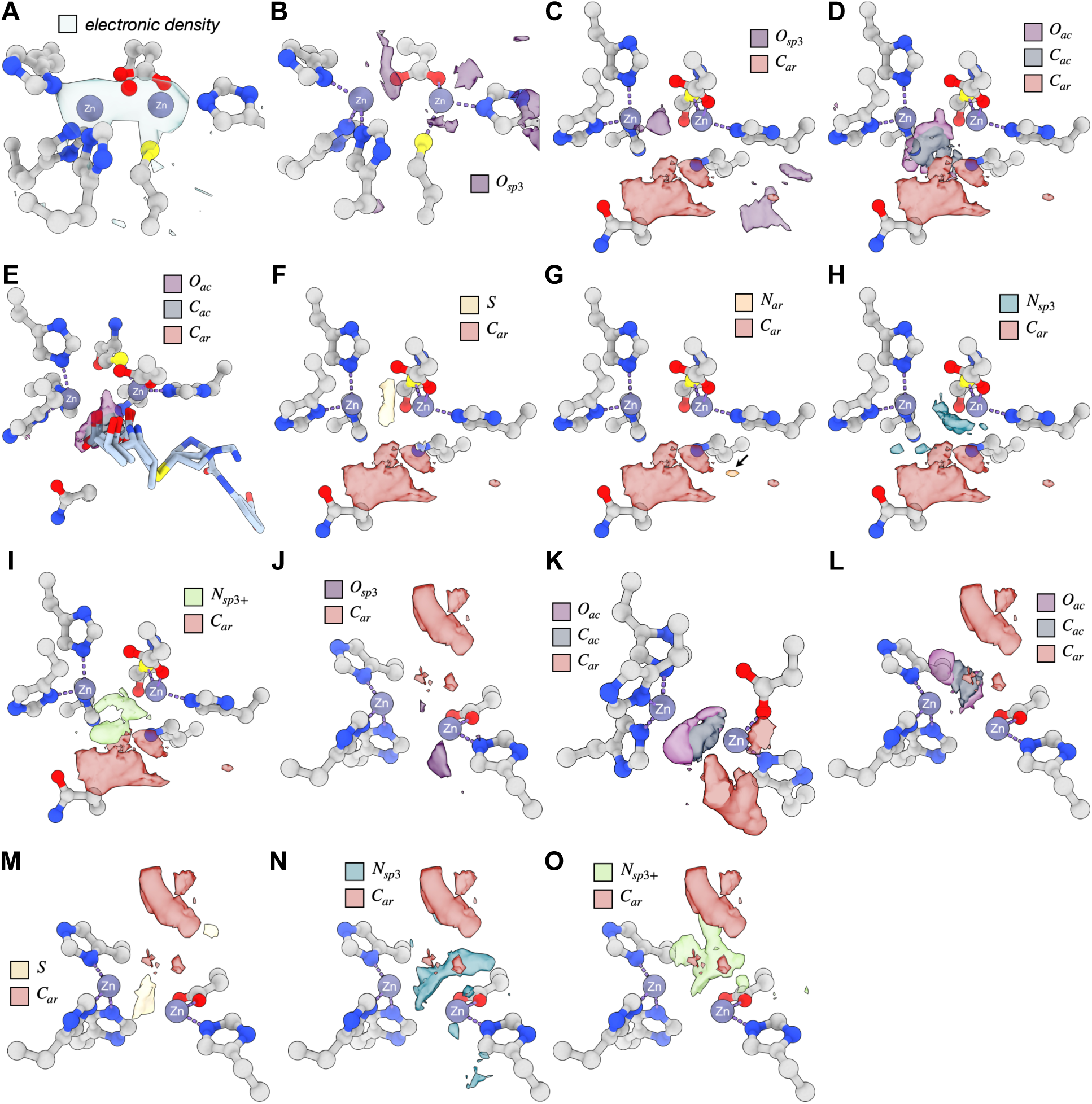
TPM Predictions for the IMP13 Metalloenzyme. A) The experimental electronic density of apo IMP13 (PDB-ID 6R78) showing the presence of an oxygen atom between the zinc cations (hydroxy group).^31^ B-I) TPM predictions for IMP13 with hydrolyzed meropenem (PDB-ID 6R73). B) sp^3^ oxygen signals overlay with good accuracy with the bridging oxygen atom between the zinc atoms, even though the structure used to generate TPMs was not of the apo protein. C) Alternative view of the predictions in the previous block, showing the relative position of oxygen atoms between the zinc ions. Inclusion of aromatic signals. D) Prediction of carboxyl groups and E) overlay of those signals with the structures of several inhibitors containing the carboxyl group. F) Predicted sulfur clouds. G) Predicted aromatic nitrogen clouds. Although aromatic nitrogens and sulfur atoms share some properties, e.g., binding to metal ions, the predictions are atom-type specific. On the other hand, H) amine nitrogens are sought between the zinc cations. I) The TPM prediction for ammonium groups is more scattered. Although these atom types are incompatible with zinc cations, they share similar chemical properties. J-O) Similar predictions for IMP13 with hydrolyzed doripenem (PDB-ID 6S0H).

### Prospective Medicinal Chemistry Validation of TPM-Guided Design

Comparison of TPM predictions with known ligand placements in binding pockets can generate hypotheses for improved binders. However, retrospective analyses are inherently susceptible to interpretive bias. We therefore tested this hypothesis prospectively by synthesizing molecules targeting a difficult protein-protein interaction interface. We chose the protist<u>s’</u> PEX14-PEX5 interface for medicinal chemistry development because previous optimization efforts had proven challenging (**Figure 5A**).^38,39^ Prior optimization appeared to have reached a plateau: conventional structure-based design strategies and medicinal chemistry iterations did not yield further improvements in pharmacological activity beyond the lead series (EC_50_ [T. cruzi] = 0.851 μM). To explore additional design options, we generated TPMs from the PDB ID 5L87 and overlaid them with the docking pose of lead compound **1** (**Figure 5B**). This analysis suggested two structural modifications that had not been prioritized previously (**Figures 5B, C**): 1) on the left side of the binding pocket, a C_sp3_ density patch suggested that introducing an alkyl substituent onto the western aromatic moiety (Ar^W^) might improve local complementarity; and 2) examination of the C_ar_ density indicated that attachment of the eastern moiety (Ar^E^) to the central scaffold was suboptimal, because substantial portions of this group did not align with TPM features. A one-atom shift in the attachment point of Ar^E^ to the central scaffold, yielding a regioisomer, was therefore predicted to improve alignment. TPMs also showed an N_sp3+_ density patch near the negatively charged residues Glu34 and Asp38, consistent with the potential for hydrogen-bonding or salt-bridge interactions. This interpretation is consistent with our previous structure-activity relationship studies, in which introduction of an ethylammonium group onto the pyrazolo[4,3-c]pyridine core of PEX14 inhibitors enhanced activity.^38,39^ Because the ligand co-crystallized with PEX14 in PDB structure 5L87 does not contain this functionality, the corresponding TPM feature is consistent with learning beyond direct reproduction of the bound ligand and transferability. Guided by these TPM-derived hypotheses, we synthesized four new inhibitors, compounds **2**-**5** (**Figure 5D**), and assessed their *in vitro* efficacy against *T. cruzi*. Most TPM-guided modifications were associated with improved potency. Compound **3**, in which the chloro substituent of lead compound **1** was replaced with a cyclopropyl group, reduced parasitic growth with an approximately fivefold improvement in activity and also showed improved selectivity: the selectivity index (SI = EC_50_ [*T. cruzi*]/CC_50_ in L6 skeletal muscle cells) increased from 13.4 to 42.1. By contrast, this effect was not observed for compound **2**, which contained a methyl group in place of the chloro substituent. Compound **2** nevertheless displayed cellular activity similar to that of lead compound **1** (EC_50_ [T. cruzi] = 0.834 µM). This result is consistent with the TPM predictions, because the methyl group does not extend sufficiently to overlap with the predicted C_sp3_ density.

**Figure 5:**
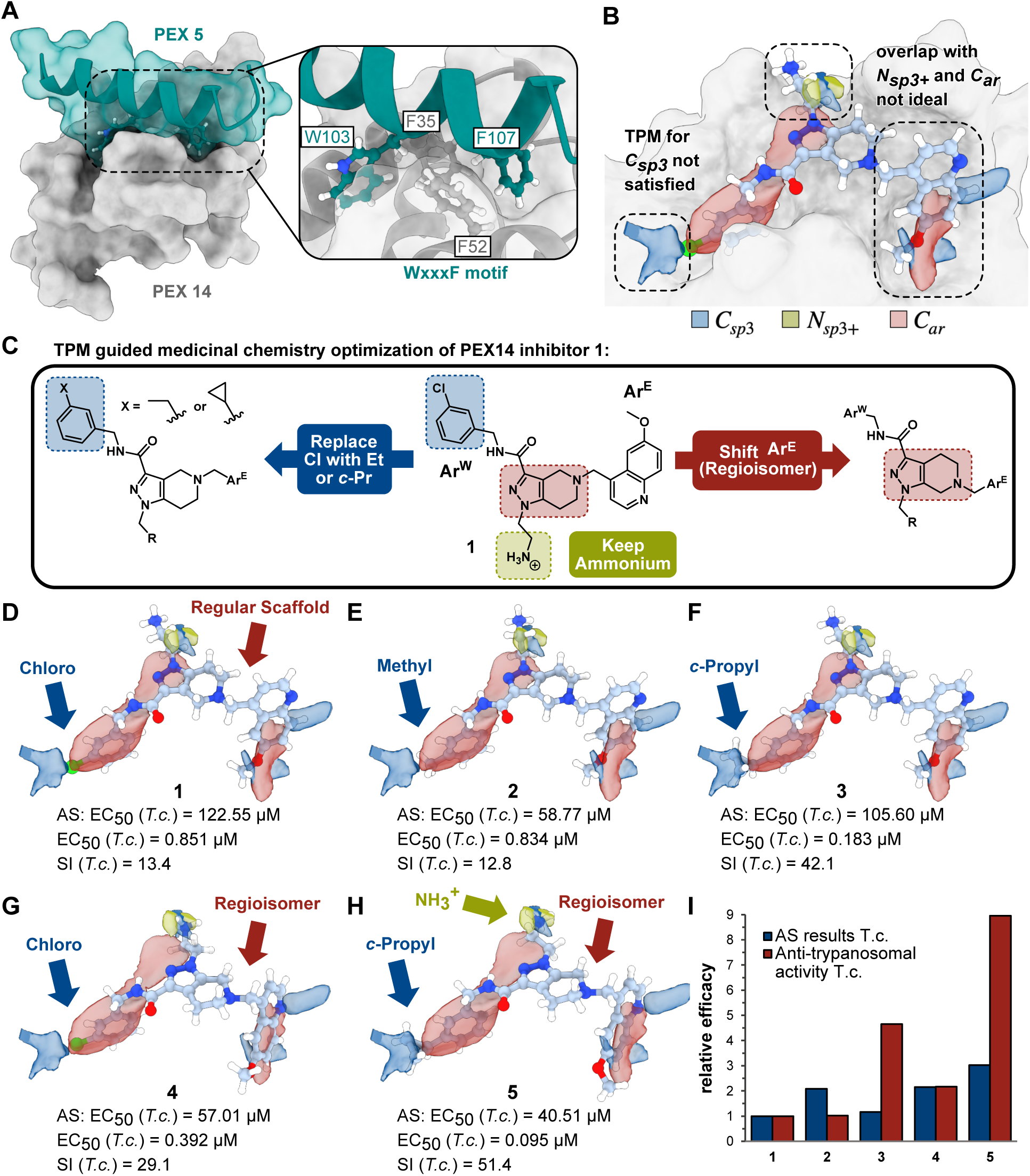
Prospective Validation of the TPM Model on a Challenging Protein-Protein Interaction (PPI) Target. A) The PPI interface of PEX14 and PEX5, and the main residues involved in the interaction. B) TPM predictions based on the 5L87 crystal structure, containing PEX14 bound to one inhibitor.^38^ The maps reveal three possible structural modifications: i) introduction of alkyl groups on the leftmost aromatic substituent (also termed western aromatic moiety); ii) shift of the rightmost aromatic group (also termed eastern aromatic moiety) to match the aromatic density prediction; iii) and introducing an ethyl ammonium group in the center of the ligand to capture the negatively charged residues Glu34 and Asp38. Note that the ligand in the reference crystal structure does not contain the ammonium group. C) Medicinal chemistry strategy used for preparing the new inhibitors. D-H) The structure and efficacy of several inhibitors, compounds **1**-**5**, along with their target selectivity. I) The relative efficacy of all compounds herein studied.

Modifying the core scaffold of a lead compound is synthetically demanding because it can require substantial changes to established synthetic routes. Accordingly, such changes benefit from strong structural justification before synthetic effort is invested. Guided by the TPMs, we generated regioisomers that showed a consistent increase in activity. Regioisomer **4** showed an approximately twofold improvement in pharmacological potency relative to inhibitor **1** while displaying improved alignment with the TPMs. This conformational adjustment also places the ligand’s ammonium group in closer agreement with the predicted N_sp3+_ density located between the two negatively charged residues. In this position, an ammonium group is consistent with potential hydrogen-bonding and salt-bridge interactions with Glu34 and Asp38. Regioisomerization is also expected to improve alignment of compound 4’s quinoline moiety (Ar^E^) with the C_ar_ TPM relative to derivatives containing the original pyrazolo[4,3-c]pyridine scaffold (compounds **1**-**3**). Finally, target molecule **5**, which incorporates the combined TPM-guided modifications and shows the strongest overall overlap with the generated TPMs, emerged as the most potent inhibitor across the biological assays, with an approximately tenfold improvement in trypanocidal activity relative to lead compound **1**. These findings support the practical value of TPM-guided hypotheses in prospective drug design. More broadly, the results suggest that TPMs may help prioritize design hypotheses that are not immediately apparent from conventional optimization trajectories. We also performed a quantum-mechanical analysis of the new inhibitor series to examine whether the observed activity trends were consistent with an independent physical interpretation. As detailed in Figure S5, these calculations were consistent with the TPM-guided hypotheses and the experimental activity trends, providing independent support for the proposed interpretation.

### TPM-based screening

A popular task in structure-based drug discovery studies is conducting primary *in silico* screening. Though economical from a financial standpoint, it comes at the cost of increased false-positive rates. We tested whether docking to the TPM abstractions is a viable alternative to conventional methods. Conformers of the Chemspace in-stock library of 5 million molecules were generated, then docked to the DYRK1A kinase as an interesting target in regenerative diabetes therapy.^40^ Since kinase selectivity plays a critical role in drug development, we simultaneously used the structurally similar GSK3β kinase as an anti-target (to which binding is not desired).^41^ Compounds were docked using stochastic global optimization^42^ combined with local gradient-based refinement to the TPMs. We decided to test the top-ranked (high DYRK1A score and low GSK3β score) compounds without human evaluation, so the selected chemistry would not be subject to human bias. The 38 selected compounds progressed toward experimental evaluation. Differential scanning fluorimetry (DSF) was used as a rapid orthogonal assay to assess compound-induced thermal stabilization of the target and off-target kinases (**Figure 6A, B**), providing biophysical support for selectivity profiling. Several compounds produced a measurable stabilization of DYRK1A, with six reaching at least 50% of the thermal shift observed for the reference compound harmine.^43^ By contrast, only two compounds produced a 20-30% shift for GSK3β relative to the control, whereas the remainder remained below 20%. While DSF was used to indirectly monitor target engagement via thermal stabilization, kinase activity inhibition was assessed using the ADP-Glo assay.^44^ **Figure 6C** shows broad agreement between the DSF-based ranking and the functional activity measurements. We then tested the best-scoring AI6, AI26, and AI37 molecules for their influence on MIN6 insulinoma cells and *ex vivo* isolated mouse islets (**Figures 6D-F**). Surprisingly, the compounds performed better than the reference compound harmine, which was tested in clinical trials (NCT05526430) as a curative approach in diabetes.^45,46^

**Figure 6:**
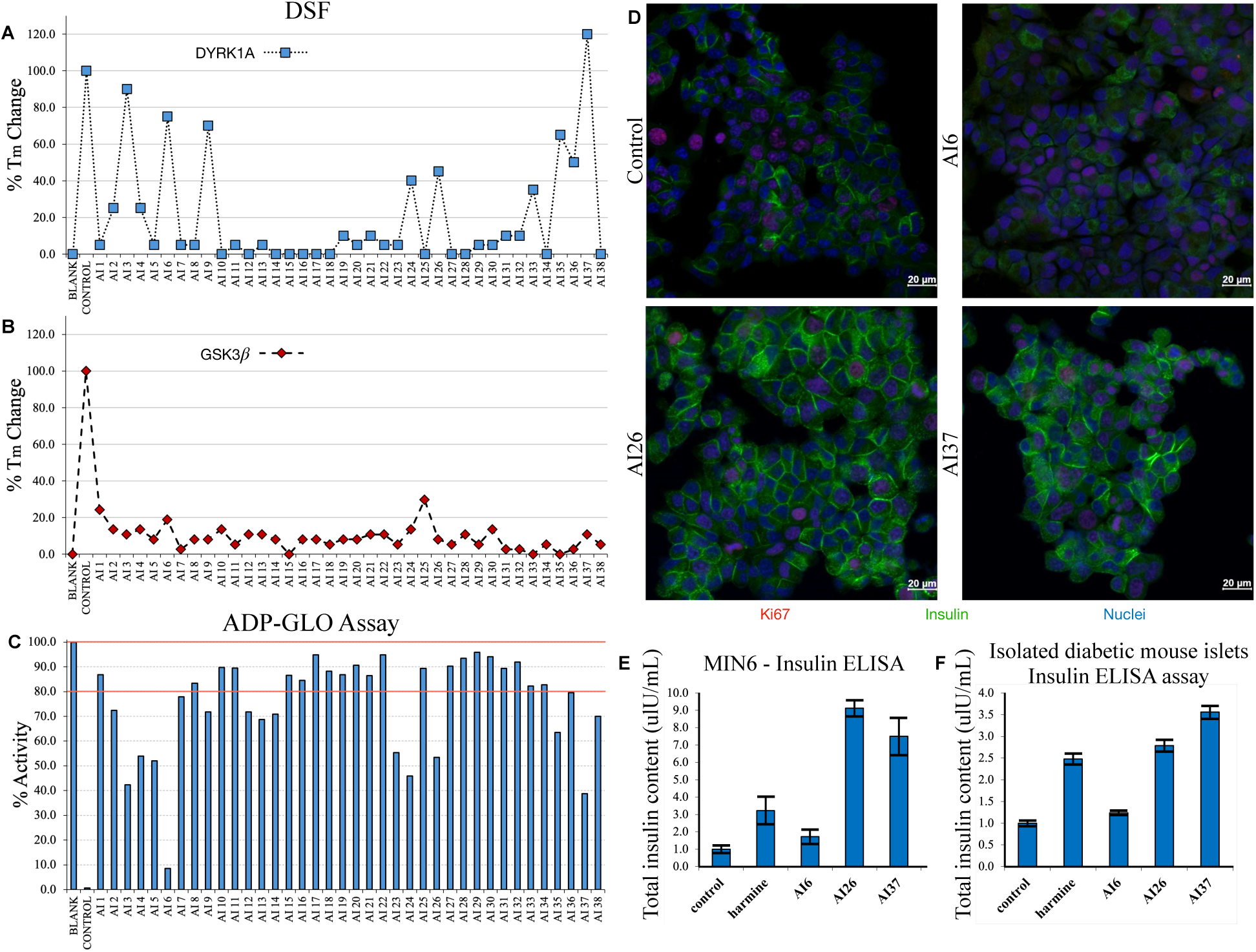
Prospective Evaluation of TPM-Guided Prioritization for Kinase-Selective Inhibitor Candidates. Compounds were prioritized to maximize their agreement with DYRK1A TPMs while minimizing overlap with GSK3β TPMs. Dye-based Thermal Shift Assay comparing thermal stabilization of the target DYRK1A kinase A) against the structurally similar B) GSK-3β. The compounds increase the melting temperature of the target DYRK1A kinase but have little effect on the structurally similar anti-target, indicating that TPM-based selection could identify active compounds based on their selectivity profiles. C) Enzymatic inhibition measured by the ADP-GLO kinase assay on the 38 top-scored TPM matches. The top-scoring compounds were assayed without human selection. Most of the purchased compounds show a significant reduction in DYRK1A kinase inhibition. The strongest inhibitors were assayed in cellular and isolated diabetic mouse models. D-F) Insulin (INS) and proliferation index (Ki67) in MIN6 cells were assessed before and after treatment with AI6, AI26, and AI37. The compounds promote insulin secretion in a D-E) glucose-stimulated insulin secretion (GSIS) assay and F) in isolated murine pancreatic islets. Scale bars at 20 μm. The insulin ELISA assay was performed by incubating cells/islets with the selected inhibitors. Values are normalized to the total amount of insulin, and insulin secretion is expressed as a fold increase where the basal level is set at 1. Data are expressed as mean ± SEM, N=3; *p < 0.05, **p < 0.01; ***p < 0.001 as compared to the control.

## Discussion

We demonstrated that an ML model can learn chemical- and environment-specific local interaction preferences from experimental structural data. The model properly “discovers” principles of non-bonding chemistry beyond simple two-body interactions, such as H-bonds or π-stacking, without prior physical conditioning. Using experimentally observed local atom-level environments in drug-receptor complexes, TPMs estimate the likelihood of different atom types at positions within the receptor’s pocket. By expressing predictions as non-mutually exclusive, spatially resolved atom-type probability-density maps, the model outputs a directly interpretable visual description of the “ideal” drug based solely on receptor structure. The sensitivity of TPMs to local structural details also suggests that they may be useful for comparing interaction preferences across closely related receptor states, including those induced by point mutations.

Biological and chemical diversity, coupled with limited and historically biased training data, have hampered, so far, the ability of even the most advanced models to generalize beyond what evolution and research have already observed.^1,3,6^ The high-capacity models risk dependence on ligand-level memorization, particularly when evaluated on molecules or targets that differ substantially from the training set.^1,2^ To mitigate this, we shifted the focus from whole molecules to local atomic microenvironments. To address this problem, we propose a reductionist approach, compartmentalizing drug-receptor training space into over 1.1 million local atom-level training instances derived from protein-ligand structures. In this way, we made the model agnostic to global drug and receptor features (chemotype, fold, *etc.*). Inference obtained in this way proved to be chemically meaningful and experimentally useful across several typical drug-discovery tasks.

By excluding ligand topology from the input representation, TPMs reduce dependence on whole-ligand connectivity and may lessen the bias arising from historically overrepresented regions of the chemical space, which also influences docking and scoring algorithms. TPMs shift non-bonded interaction interpretations from classical physics-grounded, 2-body intermolecular interactions (force fields) towards probabilistic representations. While not trained on quantum chemical data *per se*, the model’s inference clearly indicates its ability to rationalize beyond classical 2-body, Newtonian interpretation.

Docking against TPM-derived receptor preferences, although performed as a simple proof-of-principle experiment, has yielded potent, pharmaceutically attractive molecules with an equivalent activity profile to a clinical candidate for the same indication. Moreover, this was achieved without human intervention. In our example, we exemplified how the non-trivial problem of drug selectivity can be tackled using TPM abstractions from target and anti-target structures.

In ML settings, TPMs may also serve as conditioning signals for diffusion-based molecular generators. In principle, the TPM approach can also be used to learn protein-protein interactions. In such a case, the limited chemical diversity of biomolecules suggests re-designing (simplifying) the embeddings. They may further be combined with molecular dynamics or quantum-chemical analyses in future workflows, although such applications remain to be explored. Overall, TPMs provide an interpretable representation of local binding preferences that may complement existing computational drug-discovery methods and workflows.

TPMs can be visualized in standard structural biology software using the MRC format, facilitating practical inspection of the maps and easy interpretation by structural biologists and chemists. Their visual similarity to electron density was designed to facilitate human interpretation. In its simplistic appearance, TPM clouds can be compared to classical pharmacophores. However, the main difference is that pharmacophores are derived by applying a basic physical model to molecules, whereas TPMs are ML generalizations that impose no physical constraints. Pharmacophores usually describe discrete, local features, not continuous probability functions that cover entire interaction areas, as TPMs do.

The proposed approach intentionally excludes affinity data, which are often noisy, assay-dependent, and difficult to separate from ligand-specific context.^47–49^ Instead, we treat the observation of a ligand atom in an experimental structure as evidence that the corresponding local environment is compatible with binding. This representation inevitably introduces noise, because some observed atoms may contribute only weakly to binding; resolving this more precisely would require atom-specific energetic decomposition.^25^

### Limitations

While TPMs provide an interpretable framework for learning non-bonded interaction preferences, several limitations should be noted. The current TPM model is trained exclusively on static, experimentally resolved structures and therefore does not capture receptor dynamics, allosteric effects, or induced fit beyond what comes from observing of the same receptor region in different experimental structures. As a result, conformational ensembles and entropic contributions remain outside the model’s scope. This limitation is less severe when the receptor structure is already pre-organized by a bound ligand or cofactor (*e.g.*, ATP), but it may be more important for apo structures captured in solvent-adapted conformations. Future versions could address this by incorporating molecular-dynamics-derived structural ensembles.^25^ In addition, the model operates at the level of local atomic microenvironments and intentionally excludes explicit chemical connectivity. Although this abstraction may support generalization, it also omits intramolecular factors such as strain, torsion, and ligand conformational energetics, which are often important during compound optimization. Affinity information is likewise excluded from training to avoid conflating structural inference with noisy or context-dependent potency data. While this choice supports the identification of preferred local interactions, it limits the model’s direct applicability to predicting absolute binding free energies or ranking closely related analogs. TPMs should therefore be viewed as interaction-preference and hypothesis-generation models rather than as general affinity predictors. The training data, although diverse, also retain historical chemical and structural biases inherent to PDB-deposited complexes. E.g. some atom classes are represented in minimal numbers or not represented at all. Finally, the current implementation does not explicitly incorporate physical terms such as electrostatics, dispersion, or solvation, which could improve performance or interpretation in specific edge cases. Addressing these limitations by incorporating structural ensembles, energetic modeling, or multimodal learning could further broaden the applicability of the TPM framework.

### Related Work

Gainza *et al.* applied geometric deep learning to protein molecular surfaces represented as meshes to learn geometric and chemical features relevant to biomolecular recognition.^50^ This line of work was further extended to experimentally validated design settings, including de novo protein design^51^ and protein-switch engineering.^52^ Marchand *et al.* additionally showed that structural features learned from natural protein interfaces can generalize to previously unseen “neosurfaces” formed by proteins and small-molecule ligands.^52^

Related developments in small-molecule design have focused on generating ligands conditioned on three-dimensional pocket information. Schneuing *et al.* proposed a geometric diffusion model for ligand generation from protein pocket geometry,^53^ while Sverrisson *et al.* used point-cloud-based structural representations.^54^ Other approaches have employed discrete voxel grids, as in the work of Pinheiro *et al.*,^55^ or implicit neural fields, as in the score-based framework introduced by Kirchmeyer *et al.*^56^ These methods are related to TPMs in their use of three-dimensional structural representations, but they are primarily aimed at ligand generation rather than at predicting atom-type-resolved spatial preference maps.

TPMs are also related in spirit to earlier hotspot, pharmacophore, and knowledge-based interaction mapping approaches, which likewise represent spatial binding preferences within protein pockets. In particular, knowledge-based statistical potentials such as DrugScore^57^ derive interaction preferences from experimental structural data and have been used for scoring and pose evaluation. TPMs differ from these approaches in three main respects: they are trained on local atom-level environments rather than on ligand-level scoring targets, they exclude ligand topology from the input representation, and they predict continuous atom-type-specific occupancy preferences intended to support interpretation and ligand optimization rather than absolute scoring. In this sense, TPMs are best viewed as an interpretable interaction-preference model positioned between classical spatial interaction mapping and modern structure-conditioned machine-learning approaches.

## Methods

### Dataset Construction

Protein-ligand complexes were obtained from the PDBBind database.^26^ The corresponding protein sequences were extracted from the PDB sequence reference file (pdb_seqres.txt) using a custom preprocessing script, yielding a curated FASTA file containing sequences for the PDBBind entries only. To reduce redundancy and ensure sequence-level independence between the training and test data, protein sequences were clustered with CD-HIT^1^ at a 70% sequence identity threshold. This procedure resulted in 4,294 non-overlapping sequence clusters. For the construction of training and test sets, a cluster-stratified splitting strategy was employed. Clusters were randomly permuted with a fixed random seed to ensure reproducibility, and 15% of clusters (approximately 654 clusters, corresponding to 4,944 protein-ligand complexes) were assigned to the test set. The remaining 85% (approximately 3,640 clusters, 14,508 complexes) were allocated to the training set. By enforcing separation at the cluster level, homologous proteins were restricted to a single partition, thereby enabling a stringent evaluation of model generalization to previously unseen, structurally distinct targets.

### Model Construction

To predict the spatial localization of ligand atoms within protein binding sites, we developed a Transformer-based neural network (∼230,000 parameters) that estimates, for any point in a pocket, the confidence with which a ligand heavy atom of a specified type occupies that location. The model architecture consists of three stacked self-attention blocks, followed by a one-vs-all classification head that outputs atom-type-specific occupancy confidence scores. Each query point is characterized by the 25 nearest heavy atoms from the protein structure. These atoms are one-hot encoded into 21 protein atom classes, defined by combinations of PDB atom names and residue identities. Atom type classes were selected to reflect common environments in drug-like molecules, with rare or minimally represented types (*e.g.*, silicon) omitted. Ligand atom types were assigned using Open Babel conventions,^59^ ensuring consistency with the input featurization of protein atoms. To ensure rotational invariance, we define a local coordinate system at each query point.^36^ The x-axis points toward the nearest protein atom; the y-axis is constructed by Gram-Schmidt orthonormalization of the vector to the nearest reactive (non-aliphatic) atom, constrained to be at least 30° from the x-axis; and the z-axis completes the right-handed frame. The positions of the 25 surrounding protein atoms are then expressed in this local frame and concatenated with their atom-type embeddings, producing a 64-dimensional representation per atom (32 from spatial coordinates, 32 from atom-type embedding). The combined spatial and chemical representations are processed by three self-attention blocks (four heads, key dimension 64, dropout 0.2). A global mean pooling layer summarizes the contextualized representation of the local environment, which is then passed to a feedforward classifier to yield confidence scores for each ligand atom type.

### Model Training

Training and evaluation data were derived from the PDBBind2020 dataset.^26^ To prevent information leakage between the training and test sets, particularly given the highly informative spatial arrangement of 25 receptor atoms, we clustered all receptor sequences at 70% identity using CD-HIT.^58^ From these clusters, 15% were randomly selected and held out to form the test set. This ensures that no closely related proteins appear in both the training and test splits. For each complex, ligand heavy atoms located within 5 Å of any protein atom were identified, resulting in approximately 921,000 training and 240,000 test samples. These atoms serve as the positive targets. To train the model to distinguish true ligand positions from plausible but incorrect locations, we generate negative samples by displacing each positive atom’s coordinates along a random direction by a distance drawn from a Gaussian distribution (mean 3.0 Å, std 1.0 Å, minimum 1.0 Å). If the resulting decoy location was at least 1.8 Å from all ligand atoms, it was accepted as a negative (NIL) sample. This distance threshold avoids generating trivial negatives with obvious steric clashes, while still challenging the model to learn meaningful spatial constraints. We employed a train-test split; the test set was used as validation data during training to monitor convergence, but no early stopping or hyperparameter tuning was performed based on these metrics.

### Training Protocol

We adopted a two-stage training strategy comprising initial pretraining on ligand occupancy and subsequent per-atom-type fine-tuning. In the first stage, the model was trained to distinguish ligand-occupied sites from unoccupied ones (NIL class). Note that we simply analyzed ligand-bound protein surfaces against unoccupied ones. Consequently, our analysis and training protocols are both unaware of any concept of pocket or subpocket, a necessary condition for generalization and for the model to operate independently of pocket definition. A binary cross-entropy loss was used to optimize the model over all available positive and NIL samples. This pretraining established general spatial and chemical features that discriminate between the presence and absence of the ligand. In the second stage, we fine-tuned the model separately for each ligand atom type with at least 100 positive examples in the training data. Each atom-type-specific classifier was initialized from the NIL model’s weights and trained in a one-vs-all setup. We employed dynamic resampling to ensure effective use of all training data while avoiding class imbalance: at the start of each training iteration, a fresh balanced subset of negative samples was drawn to match the number of positives. Using this procedure, we trained 26 atom-type classifiers out of the 52 defined types (the remaining types had insufficient training data). For evaluation and visualization, we focused on the subset of types with sufficient test-set representation, including: N_sp3_, O_sp3_, C_sp3_, Nil, B, I, S_sp3_, O_sp2_, C_ar_, S_sp2_, N_pl_, S_o2_, F, C_sp_, P, N_ar_, Cl, N_am_, Br, N_sp2+_, C_sp2_, N_sp2_, N_sp_. All models were trained using the Adam optimizer with an initial learning rate of 8×10^−4^. A cosine-annealing learning rate schedule was applied: five epochs for the NIL model and between 20 and 40 training iterations for the atom-type classifiers, depending on the number of positive samples per class. Each iteration drew a fresh balanced sample, effectively exposing the model to different subsets of the training data across the training run.

### Model Evaluation

Model performance was primarily assessed using the area under the receiver operating characteristic curve (ROC-AUC). This threshold-independent metric reflects the probability of correctly ranking a randomly chosen positive above a randomly chosen negative. We selected ROC-AUC as the principal metric because it is robust to class imbalance and does not rely on a predefined decision threshold. To ensure that the evaluation focused on meaningful atom-type discrimination, we excluded NIL (unoccupied) samples from this analysis. Each atom-type-specific classifier was therefore evaluated solely on its ability to distinguish its target atom type from all other ligand atom types in the held-out test set. For each model, predictions were computed across the entire test set, and ROC-AUC scores were calculated using only the relevant positive and negative ligand atom labels. To ensure statistical reliability, only atom types with at least thirty positive samples in the test set were included in the reported evaluation. Additional metrics (precision, recall, F1) were computed via balanced resampling: negative samples were partitioned into non-overlapping subsets equal in size to the positives, metrics were computed per subset, and the results were averaged. Full per-atom-type results are reported in the supplementary material (**Table S1**).

A more detailed model description and architecture are provided in the supplement.

### Map Generation

Following model training, atom-type classifiers can be applied to any point within a protein pocket to estimate the confidence that a ligand atom of a specific type occupies that location. By systematically evaluating these models across a three-dimensional grid, we generate high-resolution preference maps that visualize spatial binding propensities for individual atom types. To construct these maps, the binding site is discretized into a regular grid of cubic voxels, typically spaced at 0.25 Å intervals. This resolution provides a balance between spatial accuracy and computational efficiency. At each grid point, all trained atom-type classifiers are applied to produce a confidence score for each. The resulting maps encode the learned structural preferences derived from experimental protein-ligand complexes for each atom type across the binding pocket. The maps serve as interpretable, model-derived representations of chemical complementarity, highlighting regions of the pocket that are geometrically and chemically suited to specific ligand functionalities.

### Synthetic Validation

PEX14’s pocket lipophilicity stems from two phenylalanine amino acids, Phe35 and Phe52, which are crucial for the interaction with PEX5 via one of the several diaromatic WxxxF peptide motifs present in the intrinsically disordered N-terminal region of this import receptor protein. Inspired by this binding mode a series of small molecule inhibitors was designed and synthesized using the following main scaffold: two aromatic moieties (Ar^W^ and Ar^E^) are linked via a pyrazolo[4,3-*c*]pyridine heterocycle (**Figure 5D**). Through multiple iterative cycles of hit-to-lead optimization, a diverse array of aromatic moieties for Ar^W^ and Ar^E^, including substituted naphthyl, benzyl, phenyl, pyridazine, pyridine, quinoline, and indole derivatives, was systematically investigated. This comprehensive exploration led to the discovery of over 500 PEX14 inhibitors, several of which demonstrated significant enhancements over the initial screening hit. However, with compound **1** (**Figure 5D**), the optimization reached an apparent plateau, as conventional structure-based drug design strategies and traditional medicinal chemistry approaches failed to yield further improvements in its pharmacological properties (EC_50_ (*T. cruzi*) = 0.851 μM). Consequently, retaining this modification in the next generation of PEX inhibitors was imperative. However, the TPM model indicated that further optimization was possible, as one of the ammonium group of compound **1** did not achieve optimal alignment with the predicted N_sp3+_ density maps. We hypothesized that this alignment could improve for the planned regioisomer derivatives.

A detailed description of the synthetic routes is available in the supplement.

### Quantum Mechanical Calculations

To further examine the TPM-suggested ligand modifications, we performed quantum-mechanical calculations using our in-house ULYSSES software.^60^ Calculations and subsequent analyses followed protocols described previously.^61,62^

### Kinase Selectivity Profiling

To estimate selectivity across the kinome, we assembled a panel of approximately 100 kinase crystal structures from the PDB and aligned them to a common reference frame. On this basis, we generated TPMs covering each binding site. Because the structures are spatially aligned, TPMs at roughly corresponding positions directly encode local differences in binding-site properties between kinases. Each compound was first docked into the DYRK1A TPMs using a full binding-site search. The resulting pose was then used to initialize docking into every other kinase in the panel. The assumption is that if a compound binds a related kinase, it will adopt a similar binding mode given the conserved ATP-pocket fold. This allowed us to perform local refinement docking at each off-target rather than independent full docking runs, reducing the computational cost from N_kinases_×N_compounds_ independent campaigns to a single full dock per compound, followed by fast local optimizations. From the resulting score profiles, we derived a selectivity correction. The penalty combines the mean off-target docking score with a term that specifically punishes compounds whose off-target scores approach the DYRK1A score. For this, we applied a sigmoid-based proximity weight: for each off-target kinase k, a logistic function w_k_=1/(1+exp(−β(S_k_−S_0_+Δ))) quantifies how close its score S_k_ is to the DYRK1A score S_0_. Off-targets that score comparably to DYRK1A receive a weight near one, while weak binders are effectively ignored. Summing these weights yields a penalty that counts the number of kinases likely to be co-inhibited at a therapeutically relevant dose. The final ranking score combines the primary DYRK1A docking score with this selectivity penalty.

### Dye-based Thermal Shift Assay

To assess the thermal stability of the DYRK1A and GSK3β kinases in the presence of the tested compounds, melting temperatures (Tₘ) were determined using a Thermal Shift Assay (TSA) as described previously.^63^ Recombinant DYRK1A and GSK3β proteins (1.5 mg/mL) were incubated with Sypro Orange dye (1:200 dilution, Sigma-Aldrich) in 20 mM HEPES, 150 mM NaCl, 10 mM MgCl_2_, 1 mM β-mercaptoethanol, pH 8.0, and 10 µM of the compound or DMSO as a control. The fluorescence signal of Sypro Orange was measured as a function of temperature from 5 to 95 °C with 0.5 °C/min increments using excitation at 492 nm and emission at 610 nm. The melting temperature (Tₘ) was calculated as the inflection point of the fluorescence curve. Experiments were performed in triplicate.

### ADP-GLO Kinase Activity Assay

The inhibitory activity of the tested compounds against the DYRK1A kinase was determined using the ADP-Glo Kinase Assay (Promega). The synthetic peptide DYRKtide (RRRFRPASPLRGPPK, Caslo ApS) served as the substrate for DYRK1A. Tested compounds were initially prepared as 50 mM stock solutions in DMSO and then diluted with the reaction buffer (100 mM MOPS, pH 6.8, 100 mM KCl, 10 mM MgCl_2_) to achieve the desired final concentrations. A 20 µL aliquot of the kinase and 10 µL of the test compound were incubated together for 15 minutes at room temperature. The reaction was initiated by the simultaneous addition of 20 µL of a substrate mix containing ATP and DYRKtide, and was allowed to proceed for 30 minutes at room temperature. Following this, 5 µL of the reaction mixture was transferred to HTRF 96-well low-volume plates (Cisbio), and ATP consumption was measured using the ADP-Glo Kinase Assay kit (Promega) according to the manufacturer’s protocol. The final composition of the reaction mixture was 5 ng/µL DYRK1A kinase, 10 µM compound, 100 µM ATP, and 200 µM DYRKtide. Harmine (Sigma-Aldrich) was used as a reference compound. Each data point was collected in triplicate. The degree of kinase inhibition was calculated as the percentage decrease in signal relative to the control sample without the inhibitor.

## Code Availability

The source code for the TPM model and the scripts required to reproduce the main analyses are available at GitHub: link. A web server for TPM generation and analysis is currently under construction and will be available at link.

## Acknowledgements

This work was supported by the Bundesministerium für Bildung und Forschung (BMBF), projects SUPREME (number: 031L0268) and DATIPilot Sprint–MISATO (03DPS1234); by the Bundesministerium für Wirtschaft und Klimaschutz (BMWK) via ZIM (grant number: KK5710001BA4); by the Deutsche Forschungsgemeinschaft, DFG grant CRC387, project ID TRR 387/1-514894665; by the National Science Centre (NCN, Poland) research grant no. UMO-2022/45/B/NZ7/04269.

## Supplemental Figures

**Figure S1. Additional validation of the TPM model for VEGFR crystal structures with sorafenib, sunitinib, and axitinib.**

A) Overlay of sorafenib and axitinib structures and the TPM predictions for sorafenib in VEGFR. Axitinib’s pyridyl ring’s nitrogen is not involved in any interaction with the target, and the TPM model reflects this. B) TPM predictions for the crystal structure of sunitinib in VEGFR (PDB-ID 4AGD) and overlay with the binding pose of other ligands considered in this analysis. C) TPM predictions for axitinib in VEGFR (PDB-ID 4AGC) and overlay with the binding poses of other ligands considered.

**Figure S2: Ligand imprint in experimental crystal structures.**

A) Overlay of the crystal structure of imatinib in BCR::ABL1 (PDB-ID 2HYY, ligand color code white, protein color code light blue) with that of staurosporine in BCR::ABL1 (PDB-ID 2HZ4, ligand color code brown, protein color code darker blue). TPMs predicted for the latter structure reflect the ligand’s chemical composition in the pocket and differ from the TPM predictions for imatinib in BCR::ABL1 (see Figure 2). Similar overlay with dasatinib (PDB-ID 2GQG) instead of imatinib. Dasatinib inherits imatinib’s color code in this block. C) Overlay of dasatinib’s structure with the TPM predictions for the crystal 2HZ4, which contains staurosporine in BCR::ABL1. D) Staurosporine overlaid with the TPMs for 2HZ4. E) Overlay of dasatinib and staurosporine with the TPMs predicted for the crystal structure 2GQG, which contains dasatinib in BCR::ABL1. Color code identical to that of block B). F) Dasatinib overlaid with the TPMs predicted for 2GQG, the crystal structure of dasatinib in BCR::ABL1. G) Staurosporine overlaid with the TPMs predicted for 2GQG. H) TPMs for 2HZ4 (staurosporine in BCR::ABL1). The latter are to be compared against predictions for I) 2GQG (dasatinib in BCR::ABL1), and J) 2HYY (imatinib in BCR::ABL1). K) Overlay of imatinib (brown) and dasatinib (light blue) with the TPMs for the PDB 2HYY (imatinib in BCR::ABL1). L) Overlay of imatinib (brown) and dasatinib (light blue) with the TPMs for the PDB 2GQG (dasatinib in BCR::ABL1).

**Figure S3. The specificity of halogen signals in TPM predictions.**

A-C) TPM predictions for the crystal structure 3DV3, the structure of the MEK kinase with halogen-containing inhibitors. A) The bromine, B) chlorine, and C) iodine predictions agree to a good extent, although some signals are specific to iodine atoms. D-F) TPM predictions for the crystal structure 7AKB, containing DYRK1A with a halogen-containing ligand. Predictions for D) bromine and E) chlorine are almost interchangeable, but the same does not apply to F) iodine.

**Figure S4. TPM predictions for the 5-HT2A GPCR.**

A) The receptor’s structure with the ligand’s density, which might be serotonin, lysergic acid diethylamide, or its 2-bromo derivative. B-D) Overlay of B) bromine, C) chlorine, and D) aromatic carbon densities predicted from the cryoEM structure of the receptor with serotonin. E) Overlay of the above-mentioned predicted aromatic carbon clouds and the structures of serotonin and lysergic acid diethylamide. Similar overlays, but we consider F) aromatic nitrogens or G) ammonium nitrogens instead. H) Overlay of serotonin, 2-bromo lysergic acid diethyl amide and the bromine signals predicted for the GPCR with serotonin in the pocket.

**Figure S5. Quantum Mechanical validation of the TPM-generated ligands.**

A) Correlation plot between the experimentally measured IC50 and the quantum mechanical binding energy. B) The relative Energy Decomposition and Deconvolution Analysis (EDDA) plot, which shows that compound **5**’s improved efficacy results from a mixture of improved polarization and solvation effects. C) Electrostatics maps for the 5 ligands analyzed. All maps are similar, indicating small fine differences. D) The total interaction maps for the ligand series, followed by E) the polarization maps, F) overlap-repulsion, and G) solvation.

### TPMs are sensitive to subtle structural variations of the receptor

Next, we tested TPM sensitivity to fine structural differences between two structurally similar disease-associated kinases, GSK3β and DYRK1A (accession codes 7Z5N, 7Z1F, and 7Z1G; Figure S6). To minimize the possible drug-induced fit bias, we selected structures with the same inhibitor in the pocket. The TPM signals show resemblance for all three cases. For instance, the TPM densities for aromatic nitrogen atoms (N_ar_; **Figures S6A, B**) seem to track the hinge residue to which they are hydrogen-bonded. However, maps for aromatic and sp^3^ carbons (C_ar_, **Figure S6C**; C_sp3_, **Figure S6D**) show different shapes and extents, while maintaining their positions in the pocket. Contrary to classical pharmacophore approaches, which do not contain sp^3^-like density predictions, the model is, in this case, predicting the idealized placement of a ligand’s scaffold to optimize binding. To test whether these subtle differences result from differing atomic coordinates of pocket residues, we mutated DYRK1A’s Met240 to Gly (mutation 1) to open the binding site and to Tyr (mutation 2), to mimic GSK3β (**Figures S6E**-**G**). Expectedly, N_ar_ signals remained unchanged (**Figure S6E**), as these modifications cannot affect the hydrogen-bonding properties of DYRK1A’s hinge region. Yet, the Gly240 mutation allows the TPM-C_ar_ cloud to expand (**Figure S6G**). Introducing a Tyr side chain, on the other hand, constricts the extent of the aromatic cloud (**Figure S6G**). This example shows that the TPMs are sensitive to local binding-site composition and do not retain overall receptor-drug elements.

Next, we examined TPM sensitivity to subtle structural differences between two closely related disease-associated kinases, GSK3β and DYRK1A (PDB IDs 7Z5N, 7Z1F, and 7Z1G; Figure S6).^41^ To reduce potential bias from ligand-induced conformational changes, we selected structures containing the same inhibitor. The resulting TPM signals show substantial similarity across all three cases. For example, the TPM densities for aromatic nitrogen atoms (N_ar_; **Figures S6A, B**) track the hinge region involved in hydrogen bonding. By contrast, maps for aromatic and sp^3^ carbons (C_ar_, **Figure S6C**; C_sp3_, **Figure S6D**) differ in shape and extent while remaining localized to similar regions of the pocket. Unlike classical pharmacophore approaches, which typically do not explicitly represent sp^3^-like density features, TPMs provide a spatial description of preferred ligand scaffold placement. To test whether these differences arise from local pocket composition, we mutated DYRK1A residue Met240 to Gly (mutation 1) to enlarge the binding site and to Tyr (mutation 2) to mimic GSK3β (**Figures S6E-G**). As expected, N_ar_ signals remained largely unchanged (**Figure S6E**), consistent with the fact that these mutations do not alter the hydrogen-bonding properties of the DYRK1A hinge region. In contrast, the Gly240 mutation allows the TPM-C_ar_ density to expand, whereas introduction of a Tyr side chain restricts the extent of the aromatic density (**Figure S6G**). Together, these observations indicate that TPMs are sensitive to local binding-site composition rather than simply preserving receptor- or ligand-level patterns.

**Figure S6:**
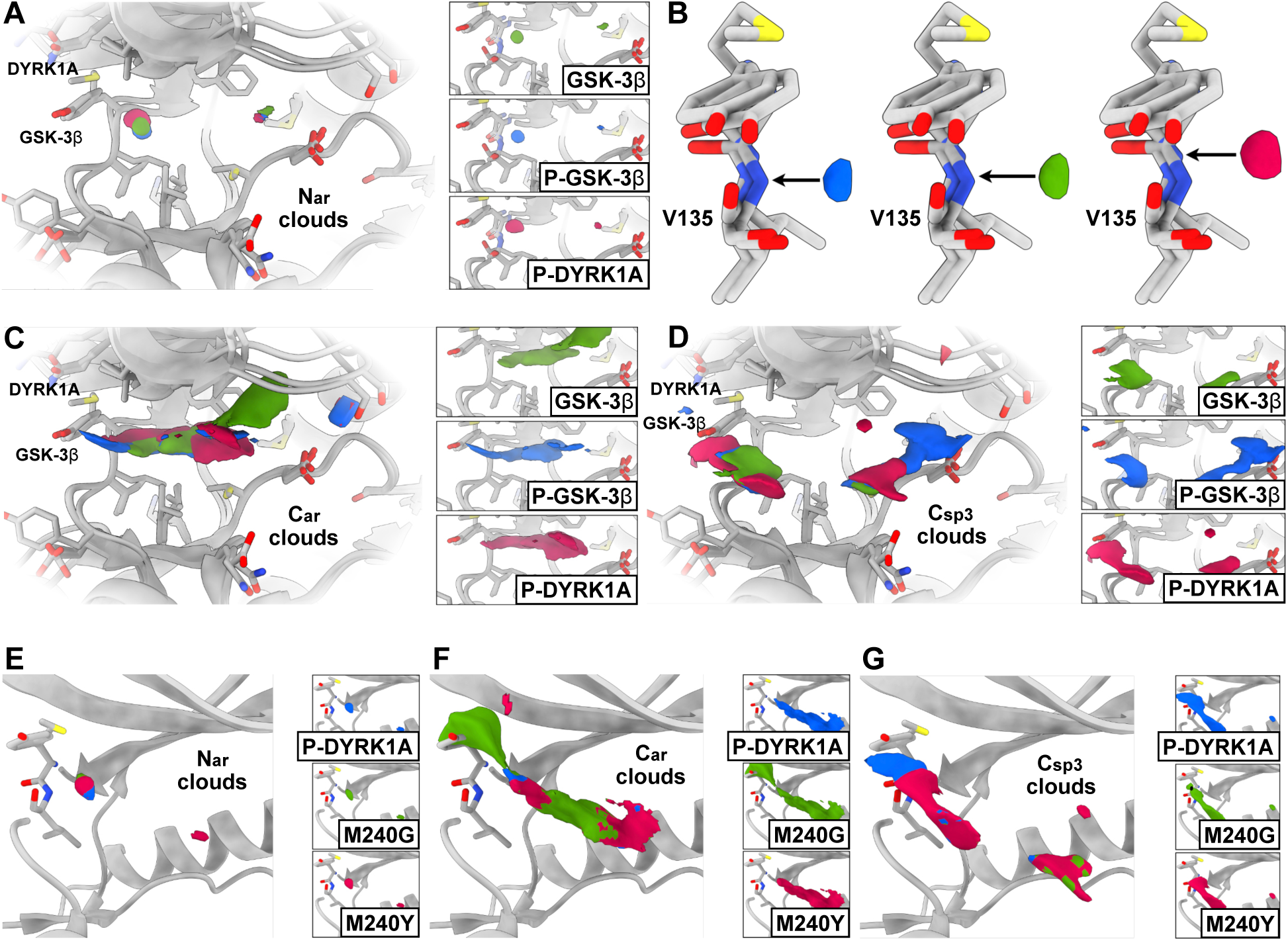
The TPM model discriminates protein pockets despite target similarity. In this analysis, we looked at TPM predictions for GSK3β and DYRK1A. A) TPMs for aromatic nitrogens follow the coordinates of hinge residues in GSK3β (phosphorylated and unphosphorylated, PDB-IDs 7Z1G and 7Z1F) and phosphorylated DYRK1A (PDB-ID 7Z5N).^41^ B) Zoom-in of the respective predictions with the kinase structures overlaid. The arrows are positioned at the geometric center of each signal. C) Aromatic carbon predictions and D) sp^3^ carbon predictions for the same systems. In both cases, cloud details are system-specific, although a general shape is somewhat conserved. Modifications in TPM-cloud predictions for mutations transforming gradually DYRK1A’s hinge into that of GSK3β. E) Although aromatic nitrogen signals remain conserved, changes are recorded for F) aromatic and G) sp^3^ carbons.

### Model Details

#### Model architecture and optimization details

Figure S7 provides a schematic of the model architecture. Briefly, the network is a Transformer-based binary classifier trained in a one-versus-all setting. The model uses pre-norm Transformer blocks with residual connections. Each Transformer block applies a layer normalization before the attention and feedforward operations. It comprises two input streams: i) distance vectors from a query point to its neighboring protein atoms, expressed in a locally defined, rotationally invariant coordinate frame constructed via Gram-Schmidt orthogonalization; and (ii) integer-encoded protein and ligand atom-type identities. Protein-atom codes are based on the convention used in the PDB format, whereas ligand atom types are grouped according to the Tripos mol2 typing used in Open Babel.^59^ The resulting geometric and chemical embeddings are concatenated into token features and processed by three identical Transformer blocks. The contextualized token representations are aggregated using 1D global average pooling and passed through a classification head comprising dropout (0.2), a dense layer with ReLU activation, another dropout (0.2), and a final dense layer followed by a softmax to produce a binary prediction. **Table S1** further summarizes model optimization details.

**Figure S7.**
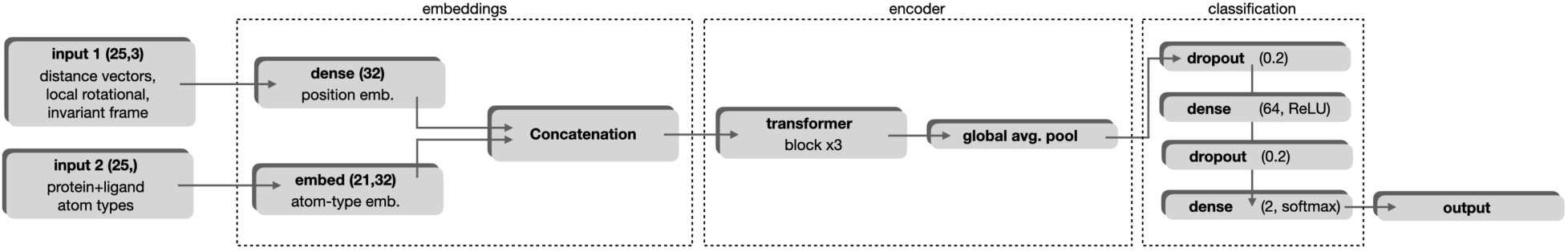
Schematic representation of the TPM model.

**Table S1:** Summary of model optimization details.

| Parameter | Value |
| --- | --- |
| optimizer | Adam |
| initial learning rate | $8 \times 10^{-4}$ |
| learning rate schedule | Cosine annealing to $1 \times 10^{-5}$ |
| batch size | 32 |
| base epochs | 20, scaled by sample count |

#### Training strategy

To train the TPM model, we follow a one-versus-all approach, defining 52 ligand atom types, of which only 26 have sufficiently large training datasets (at least 100 samples). These receive separate binary classifiers. To improve training efficiency, we use transfer learning from the NIL class (the non-ligand model) to each atom-type-specific model. For each binary classifier, training data is balanced via dynamic resampling: each iteration draws a fresh random subset of negatives matching the positive count. Further, atom-types with more samples received proportionally further training epochs. In essence, we increased the number of training epochs by one every time the log-scaled sample count doubled.

#### Data split

The TPM model was trained on the PDBbind v2020 dataset.^26^ Specifically, we used the refined set supplemented with the v2020-other-PL one. A CD-Hit,^58^ cluster-stratified split was employed for sequence clustering. To minimize any sequence-similarity leaks between the train and test sets, 15% of the stratified splits were assigned to the test set. Since the similarity-stratified analysis requires a sufficient number of test samples per bin, we opted to use the test set also for validation monitoring during training, but without early stopping or other measures that might introduce significant leakage. This was accomplished by monitoring convergence as reported by 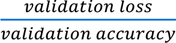 in the training logs.

#### Performance metrics

TPM metrics by category for the 26 main atom types studied in the manuscript are presented in **Table S2 below**. The mean AUC across all samples is 0.802, and the mean F1 is 0.751. AUCs for all atom types are also given in Figure S8. Logs per atom type are available in the repository attached to the submission.

**Table S2:**
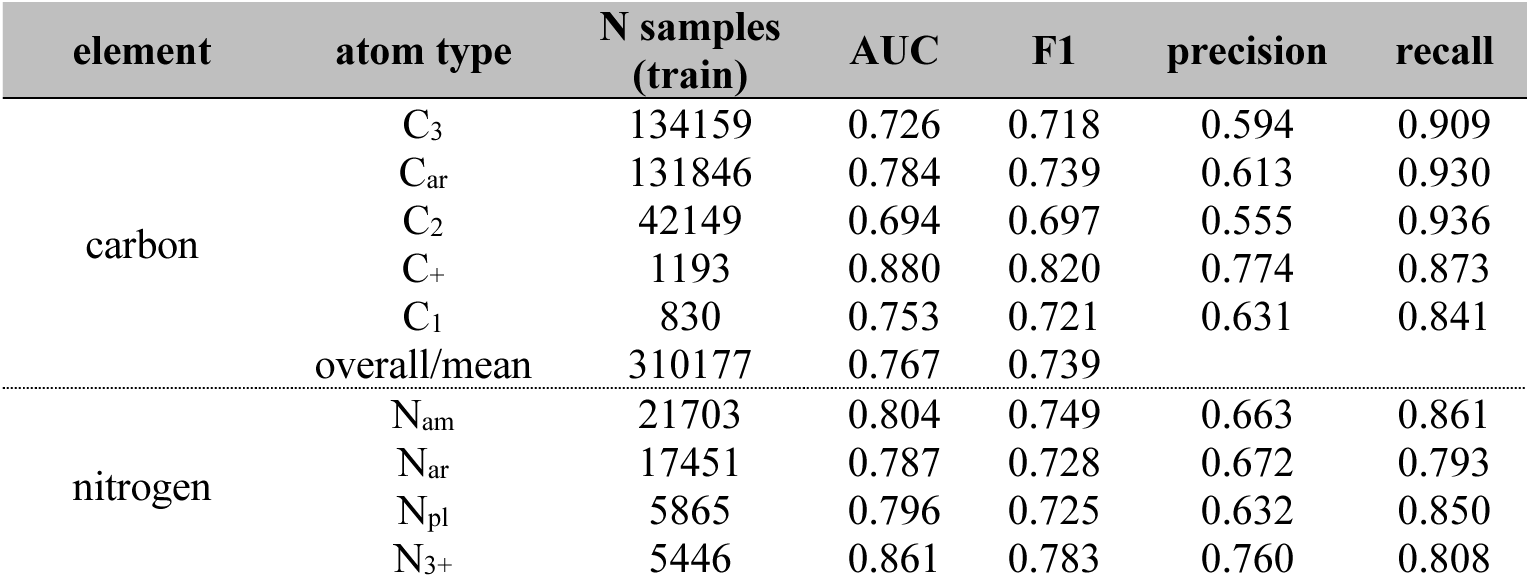
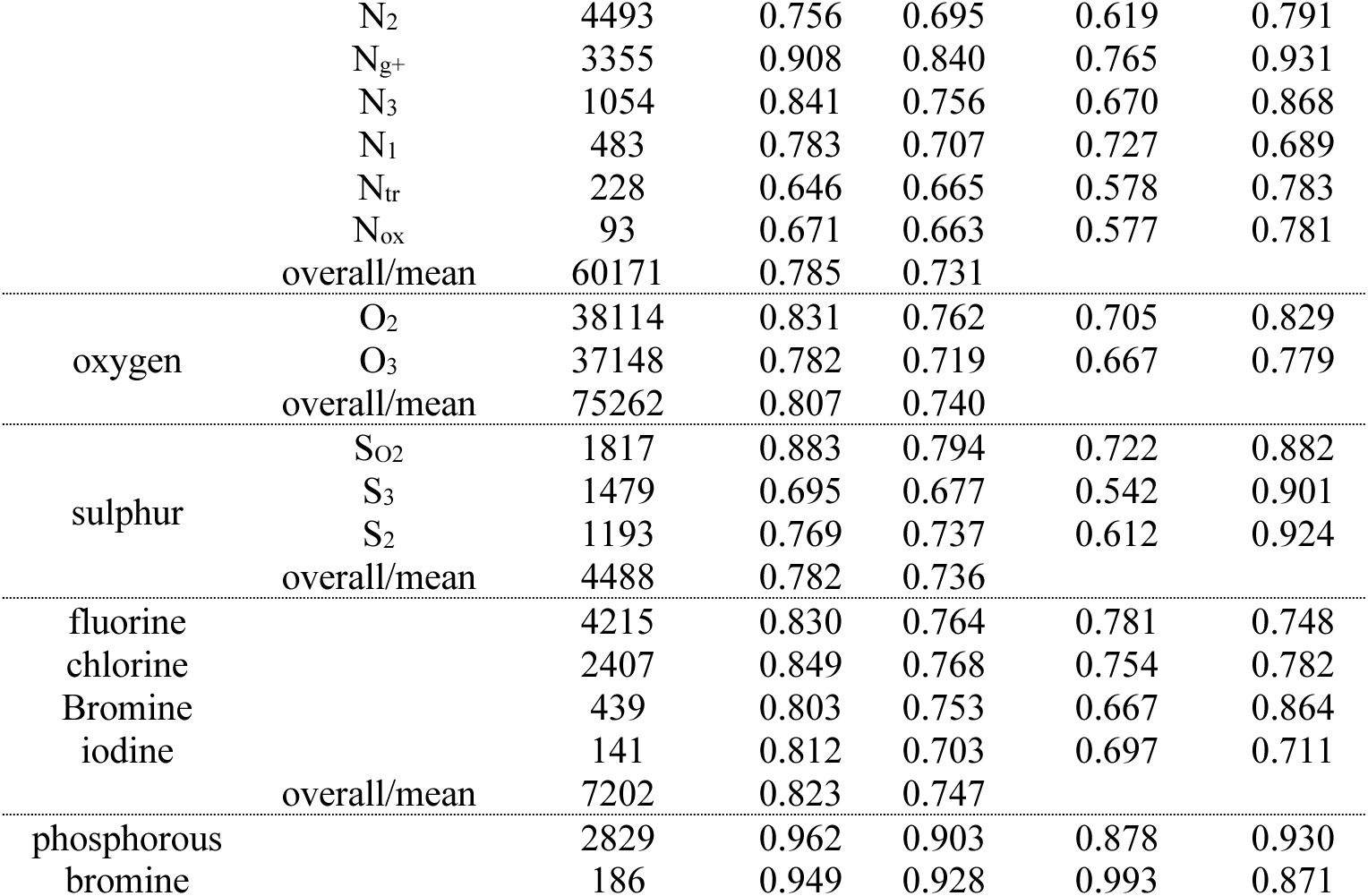
Performance metrics for the TPM model. Data is provided first for specific atom types, and then we have a summary for each category.

**Figure S8.**
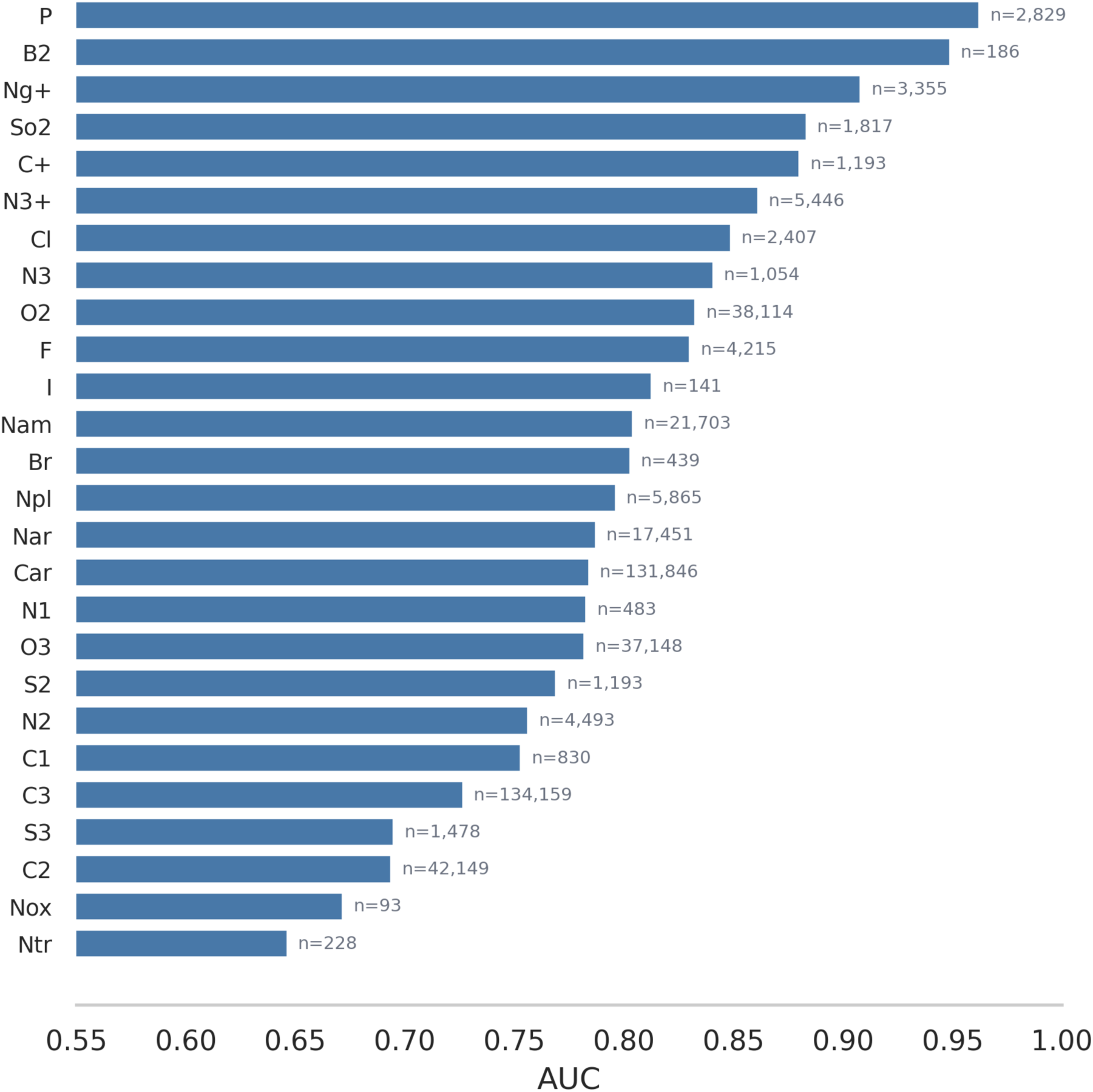
Visual representation of the AUC per atom type.

#### Learning curves

Learning curves for some of the TPM atom type channels are presented in Figure S9.

**Figure S9.**
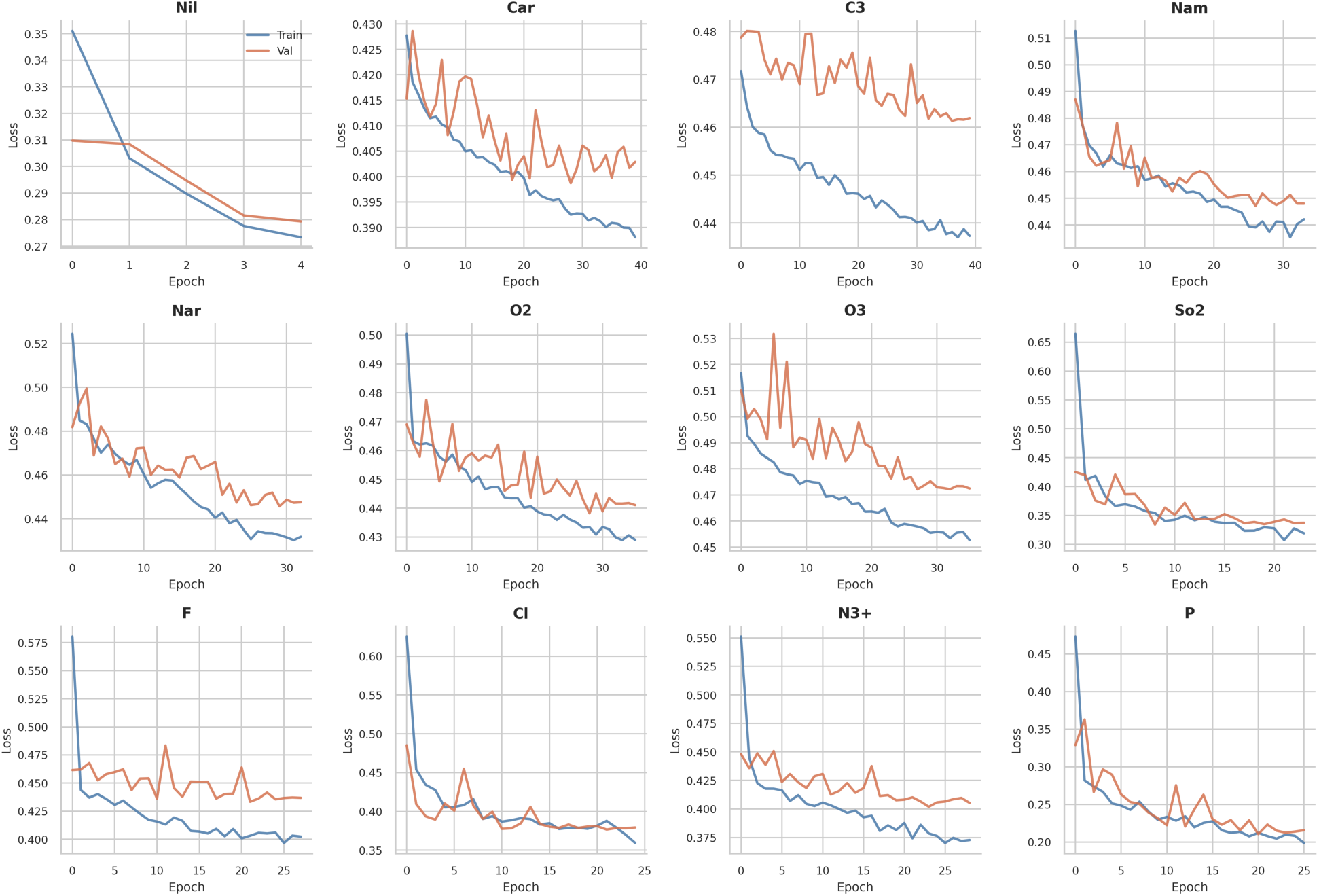
Learning curves for representative atom type channels in the TPM model.

#### Computational requirements

Training the TPM model requires NVIDIA GPUs with CUDA support. The model’s GPU memory footprint remains modestly below 1 GB at the peak. Training runs comfortably on modern CUDA-capable GPUs. The training times scale with the number of positive samples per atom type. For instance, the two largest atom type classes (C_3_ and C_ar_) require each roughly 90 minutes. Conversely, smaller classes are completed within a few minutes. In total, training all atom-type classifiers requires 7-8 hours on a single A6000 GPU.

Using the TPM model has the dependencies reported in **Table S3**.

**Table S3:** TPM model requirements.

| Package | Version |
| --- | --- |
| Python | 3.9+ |
| TensorFlow | 2.15.x |
| NumPy | 1.24+ |
| Scikit-learn | 1.3+ |
| pandas | 2.0+ |
| Matplotlib | 3.7+ |
| seaborn | 0.12+ |

### Evidence of Generalization

To address potential concerns about memorization, model performance was evaluated on stratified protein-ligand interaction (PLI) similarities to the training set using the PLINDER database.^6^ We used PLINDER because the authors specifically designed a similarity metric to detect potential train/test leakage and overfitting in structure-based models.

#### Excluding unverifiable PDBs

Test PDBs whose similarity score could not be reliably determined were excluded before stratification to prevent them from artificially inflating the “novel” (0-30 %) bin. In total, we excluded 504 structural models: either because they were not in PLINDER (192 cases due to unverifiable novelty) or because of mismatches in ligand chemical component dictionary (CCD) codes (312 cases). In the latter, the CCD mismatch case, the absence of a match in PLINDER results in PLI similarity being computed against the wrong ligand or a fallback system.

After excluding the unverifiable PDBs, we retained 3503 protein systems with a matching ligand in PLINDER and genuinely zero similarity.

#### Obtaining similarity scores

For each test PDB, the PLINDER database was queried with the *holo* flag to obtain pre-computed unique PLI query coverage metrics (uPLIqc). This required mapping each test PDB to the respective ligand CCD code using PDBbind indices. Finding PLINDER system IDs for each test PDB required exact CCD code matching wherever possible (with a part- or component-level match as a fallback). Then, for each test system, uPLIqc scores were retrieved from the *holo* database. Targets whose PDB codes appear in the training sets were filtered. The maximum similarity score was then assigned on a 0-100 % scale.

#### Stratified evaluation

After calculating similarity scores, the data were stratified according to four bins based on their similarity score to the originating PDB:

- Novel, if the similarity score lies between 0 and 30 %. This means little to no overlap with the training set.
- Low similarity, if the similarity score lies between 30 and 50 %.
- Moderate similarity, if the similarity score lies between 50 and 70 %.
- High similarity, if the similarity score is above 70 %.

Different samples from a single PDB are grouped into the same bin. Since one PDB may contribute with a manifold of samples from different bound ligands.

To get the full per-atom-type evaluation, the pipeline is rerun on the bin-specific samples. Note that, for each bin:

- NIL samples are excluded because they are non-ligand.
- Positive and negative samples are balanced by undersampling the larger class.
- The trained, atom-type-specific one-versus-all classifier is loaded to produce probability predictions.
- The AUC is computed on the full, imbalanced data.
- Precision, recall, and F1 are computed by bootstrap resampling. Negatives are partitioned into non-overlapping balanced subsets, and metrics are computed for each subset. The results are then averaged.
- Atom types with less than 20 positive samples in a bin are excluded.

This leads to bin-specific, atom-type-resolved metrics (precision, recall, F1, AUC, accuracy, and positive sample count).

#### Results by similarity bin

The main results of our stratified evaluation are shown below, in **Table S4**.

**Table S4:**
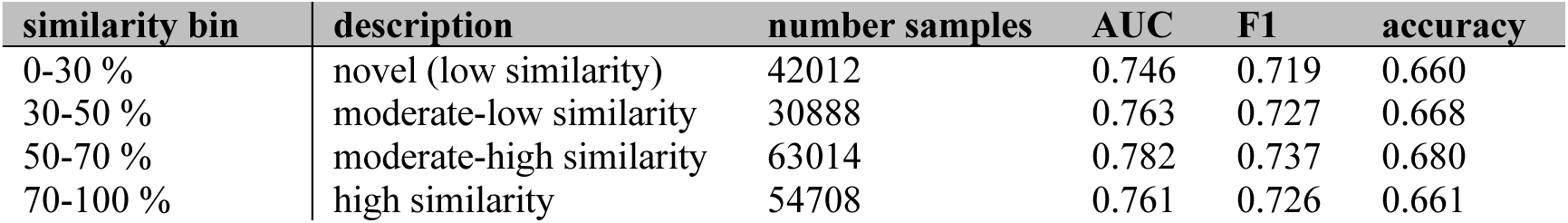
TPM model novelty evaluation.

Of the 1078 test PDBs with 0% similarity, 192 (18 %) are absent in the PLINDER database, 312 (29 %) contain a CCD code mismatch (different ligand nomenclature), and 574 (53 %) contain truly novel protein-ligand interactions. Further, it may be seen that performance on truly novel structures (0-30 % similarity) is comparable to that on high similarity (70-100 %). The AUC difference is 0.015 (0.746 vs. 0.761), with novel structures performing only slightly lower. This demonstrates that the model learned generalizable microenvironment features of the proteins rather than memorizing the training examples. If the model were simply memorizing, degraded performance on novel structures would have been expected. In fact, the moderate similarity bin (50-70 %) achieves the highest AUC score (0.782), suggesting no simple memorization gradient.

### Synthesis of PEX14 inhibitors 1-5

All target compounds were prepared using a slightly modified version of a previously described protocol used to synthesize similar PEX14 inhibitors.^38,39^ The optimized synthesis routes are displayed in **Schemes S1** and **S2**, respectively. The detailed synthesis procedures and spectroscopic data are reported in the following paragraphs.

Air and water-sensitive reactions were performed in flame-dried glassware under an argon atmosphere. Solvents used for column chromatography, extractions, and recrystallization were purchased in technical grade and distilled prior to use. Solvents used for reversed phase chromatography and HPLC-MS analyses were purchased from Thermo Fisher Scientific in HPLC-quality. Reagents and dry solvents were purchased from Sigma-Aldrich (Merck), ABCR, Alfa Aesar, Thermo Fisher Scientific, TCI and Carl Roth, and were used without further purification. Analytical thin-layer chromatography (TLC) was performed on silica-coated plates (silica gel 60 F_254_) purchased from VWR. Compounds were detected by ultraviolet (UV) irradiation at 254 or 366 nm. Manual flash column chromatography was performed using silica gel 60 (particle size: 0.040-0.063 mm) available from VWR. Automated preparative chromatography was performed on a Grace Reveleris Prep purification system using linear gradient elution and Büchi Reveleris Silica 40 μm cartridges for normal-phase and Büchi Reveleris C18 40 μm cartridges for reverse-phase separations. ^1^H and ^13^C NMR spectra were recorded at room temperature on a Bruker AV-HD400 spectrometer operating at 400 MHz. NMR peaks are reported as follows: chemical shift (*δ*) in parts per million (ppm) relative to residual non-deuterated solvent as an internal standard (CHCl_3_: *δ* (^1^H) = 7.26 ppm, *δ* (^13^C) = 77.2 ppm; DMSO: *δ* (^1^H) = 2.50 ppm, *δ* (^13^C) = 39.5 ppm), multiplicity (s = singlet, d = doublet, t = triplet, q = quartet, m = multiplet and br s = broad signal), coupling constant (Hz) and integration. HPLC-UV/MS analyses were performed on a Dionex UltiMate 3000 HPLC system coupled with a Thermo Scientific™ ISQ™ EC Single Quadrupole Mass Spectrometer, using the following method: Thermo Scientific™ Accucore™ RP-MS LC-column (2.1 x 50 mm, 2.6 μm); gradient: 5 to 95% of acetonitrile + 0.1% formic acid v/v in water + 0.1% formic acid v/v over a 5-min period; flow rate: 0.6 mL/min; UV detection at 254 nm. All target compounds exhibited a purity greater than 95%, which was determined by HPLC-UV/MS analyses.

### Synthetic procedures and spectroscopic data for PEX14 inhibitors 1-3

**Scheme S1.**
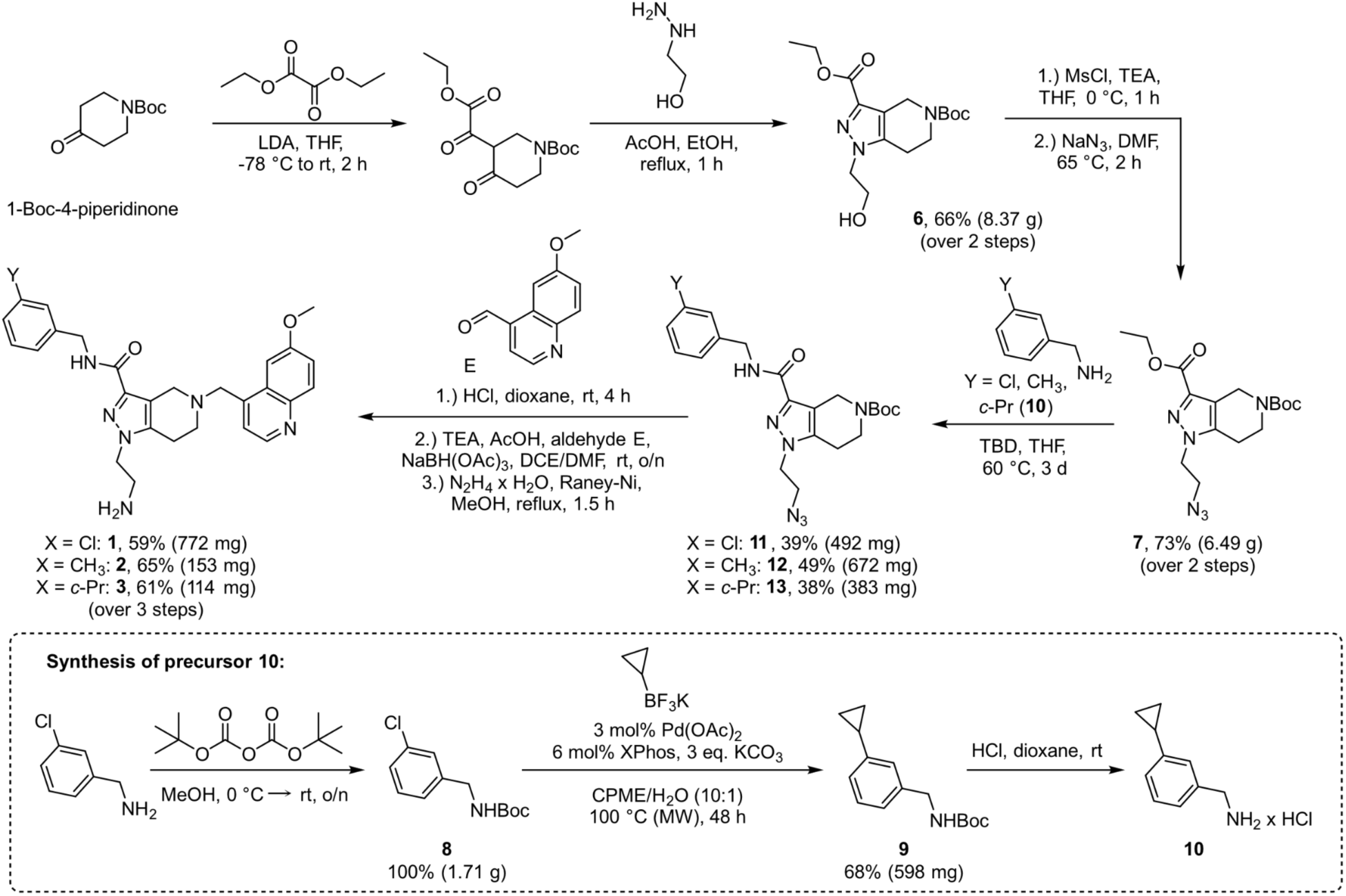
Synthetic route for the preparation of PEX14 inhibitors **1-3**.

#### 5-(*tert*-butyl) 3-ethyl 1-(2-hydroxyethyl)-1,4,6,7-tetrahydro-5*H*-pyrazolo-[4,3-*c*]pyridine-3,5-dicarboxylate (6)

A solution of lithium diisopropylamide (LDA) (2.0 M in THF, 20.6 mL, 41.2 mmol, 1.1 equiv) in dry THF (75.0 mL) was cooled to -78 °C. To this mixture, a solution of 1-Boc-4-piperidone (7.47 g, 37.5 mmol, 1.0 equiv) in dry THF (50.0 mL) was added dropwise under an argon atmosphere. After 30 min of continuous stirring at -78 °C, a solution of diethyl oxalate (6.37 g, 37.5 mmol, 1.0 equiv) in dry THF (50.0 mL) was slowly added. The resulting reaction mixture was then warmed to rt and stirred for an additional 30 min. Subsequently, 2-hydroxyethylhydrazine (2.54 mL, 37.5 mmol, 1.0 equiv), acetic acid (20.0 mL) and EtOH (25.0 mL) were added. After refluxing for 1 h, the reaction was cooled to rt and concentrated in vacuo. The remaining residue was partitioned between EtOAc (100 mL) and a saturated NaHCO_3_ solution (75.0 mL). The aqueous phase was extracted with EtOAc (3 × 50 mL) and the combined organic extracts were washed with water (50.0 mL) and brine (50.0 mL). After drying over Na_2_SO_4_, the extract was filtered and concentrated in vacuo. The obtained oily residue was then recrystallized from hexane/CH_2_Cl_2_ and subsequently dried to yield precursor **6** as an off-white solid (8.37 g, 24.6 mmol, 66%). ^1^H NMR (400 MHz, CDCl_3_): *δ* = 4.59 (s, 2H), 4.37 (q, *J* = 7.2 Hz, 2H), 4.21 – 4.18 (m, 2H), 4.03 – 4.01 (m, 2H), 3.71 (t, *J* = 5.8 Hz, 2H), 2.73 (t, *J* = 5.8 Hz, 2H), 1.47 (s, 9H), 1.37 (t, *J* = 7.2 Hz, 3H) ppm. ^13^C NMR (101 MHz, CDCl_3_): *δ* = 162.1, 154.7, 139.3, 138.2, 117.2, 80.1, 61.2, 60.7, 51.2, 28.2, 21.8, 14.2 ppm. HPLC-MS (ESI): *m/z* = 362 [M+Na]^+^; retention time: 3.30 min.

#### 5-(*tert*-butyl) 3-ethyl 1-(2-azidoethyl)-1,4,6,7-tetrahydro-5*H*-pyrazolo[4,3-*c*]pyridine-3,5-dicarboxylate (7)

Pyrazole **6** was dissolved in dry THF (50.0 mL) under an argon atmosphere and cooled to 0 °C. After addition of triethylamine (4.97 g, 6.81 mL, 49.1 mmol, 2.0 equiv) and methanesulfonyl chloride (2.66 g. 2.47 mL, 31.9 mmol, 1.3 equiv), the resulting mixture was stirred for 1 h at 0 °C. Afterwards, the reaction mixture was filtered through Celite^®^ and the solvent was removed under reduced pressure. For the second step, the crude mesylate was redissolved in dry DMF (25.0 mL) and sodium azide (4.79 g, 73.7 mmol, 3.0 equiv) was added. The mixture was then heated to 70 °C for 2 h, after which it was quenched by the addition of a saturated NaHCO_3_ solution (50.0 mL). The aqueous phase was then extracted with EtOAc (3 x 70.0 mL) and the combined organic phases were washed with water (50.0 mL) and brine (50.0 mL), dried over Na_2_SO_4_, filtered and concentrated under reduced pressure. The crude product was purified via gradient column chromatography (*c*-hexane/EtOAc: 25, 35, 50, 75, 100% EtOac), which yielded the desired product **7** as an off-white solid (6.49 g, 17.6 mmol, 73%). ^1^H NMR (400 MHz, CDCl_3_): *δ* = 4.61 (s, 2H), 4.39 (q, *J* = 7.2 Hz, 2H), 4.19 (t, *J* = 5.6 Hz, 2H), 3.78 (t, *J* = 5.6 Hz, 2H), 3.72 (t, *J* = 5.8 Hz 2H), 2.73 (t, *J* = 5.8 Hz, 2H), 1.47 (s, 9H), 1.38 (t, *J* = 7.15 Hz, 3H) ppm. ^13^C NMR (101 MHz, CDCl_3_): *δ* = 162.0, 154.7, 139.3, 139.0, 117.3, 80.1, 60.7, 60.5, 50.8, 48.3, 41.1, 40.1, 28.2, 21.8, 20.8, 14.2 ppm. HPLC-MS (ESI): *m/z* = 365 [M+H]^+^, 387 [M+Na]^+^; ^+^; retention time: 3.63 min.

#### *tert*-Butyl (3-chlorobenzyl)carbamate (8)

To a solution of 3-chlorobenzylamine (1.00 g, 7.06 mmol, 1.0 equiv) in dry MeOH (20.0 mL) Boc anhydride (1.54 g, 7.06 mmol, 1.0 equiv) dissolved in dry MeOH (10.0 mL) was added in a dropwise manner. The reaction mixture was stirred at rt overnight and the next day concentrated in vacuo. The resulting material was pure enough without any further purification and was used directly in the next step (1.71 g, 7.06 mmol, colorless oil, quantitative). ^1^H NMR (400 MHz, CDCl_3_): *δ* = 7.31 – 7.25 (m, 3H), 7.20 – 7.15 (m, 1H), 4.90 (s, 1H), 4.31 (s, 2H), 1.48 (s, 9H) ppm. ^13^C NMR (101 MHz, CDCl_3_): *δ* = 156.0, 141.3, 134.6, 130.0, 127.6 (2x C), 125.6, 80.0, 44.3, 28.6 ppm. HPLC-MS (ESI): *m/z* = 242 [M+H]^+^, retention time: 3.76 min.

Attaching the cyclopropyl group to the benzyl moiety of precursor **8** was accomplished via a Pd-catalyzed cross-coupling reaction that was reported in literature by Molander and Gormisky.^64^

#### *tert*-Butyl (3-cyclopropylbenzyl)carbamate (9)

A microwave vial was charged with Boc-protected chlorobenzylamine **8** (853 mg, 3.53 mmol, 1.0 equiv), potassium cyclopropyltrifluoroborate (549 mg, 3.71 mmol, 1.1 equiv), and K_2_CO_3_ (1.46 g, 10.59 mmol, 3.0 equiv). A mixture of CPME/H_2_O (10:1) was added as solvent, which was degassed for 30 min via an argon gas stream. XPhos (101 mg, 0.212 mmol, 6 mol%) and Pd(OAc)_2_ (23.8 mg, 0.106 mmol, 3 mol%) were added to the degassed reaction mixture. Afterwards, the tube was sealed with a cap lined with a disposable Teflon septum and heated at 100 °C for 48 h in a microwave reactor. After this time period, the reaction was cooled to rt, diluted with H_2_O (50 mL) and extracted with CH_2_Cl_2_ (4 x 50 mL). The combined organic layers were washed with brine (50 mL), dried over Na_2_SO_4_, filtered and concentrated under reduced pressure. The crude product was purified by gradient column chromatography (*c*-hexane/EtOAc: 10, 20% EtOAc) to yield the product as a colorless oil in 68% (598 mg, 2.42 mmol). ^1^H NMR (400 MHz, CDCl_3_): *δ* = 7.21 (t, *J* = 7.6 Hz, 1H), 7.05 (d, *J* = 7.7 Hz, 1H), 7.01 – 6.94 (m, 2H), 4.78 (br s, 1H), 4.27 (s, 2H), 1.88 (tt, *J* = 8.4, 5.1 Hz, 1H), 1.47 (s, 9H), 0.98 – 0.91 (m, 2H), 0.72 – 0.65 (m, 2H) ppm. ^13^C NMR (101 MHz, CDCl_3_): *δ* = 156.1, 144.6, 139.0, 128.7, 125.1, 124.8, 124.7, 79.7, 45.1, 28.6, 15.5, 9.4 ppm. HPLC-MS (ESI): *m/z* = 248 [M+H]^+^, retention time: 3.87 min.

#### 3-cyclopropylbenzylamine hydrochloride (10)

Boc-deprotection: Boc-protected 3-cyclopropylbenzylamine **9** (540 mg, 2.18 mmol, 1.0 equiv) was dissolved in dry 1,4-dioxane (15.0 mL) under an argon atmosphere. After the addition of HCl (4.0 M in 1,4-dioxane, 7.64 mL, 30.6 mmol, 14.0 equiv), the resulting reaction mixture was stirred for 4 h at rt. Afterwards, the solvent was removed under reduced pressure and the off-white residual solid was dried in vacuo. The crude product was used without further purification for the follow-up reaction step.

#### General procedure for the synthesis of precursors 11-13

A solution of pyrazole azide **7** (1.0 equiv), the respective benzylamine (1.2 or 1.0 equiv) and TBD (1.0 or 2.0 equiv) in dry THF was stirred at 60 °C for 4 d. The mixture was concentrated in vacuo and the residue was purified by gradient flash column chromatography (*c*-hexane/EtOAc: 20, 35, 50, 70, 90, 100% EtOAc).

#### *tert*-Butyl1-(2-azidoethyl)-3-((3-chlorobenzyl)carbamoyl)-1,4,6,7-tetrahydro-5*H*-pyrazolo[4,3-*c*]pyridine-5-carboxylate (11)

Pyrazole azide **7** (1.00 g, 2.75 mmol, 1.0 equiv), 3-chlorobenzylamine (402 μL, 467 mg, 3.30 mmol, 1.2 equiv), TBD (383 mg, 2.75 mmol, 1.0 equiv), dry THF (20.0 mL). Yield: 39% (492 mg, 1.07 mmol), white solid. ^1^H NMR (400 MHz, CDCl_3_): *δ* = 7.34 (s, 1H), 7.28 – 7.18 (m, 4H), 4.71 (s, 2H), 4.58 (d, *J* = 6.1 Hz, 2H), 4.13 (t, *J* = 5.7 Hz, 2H), 3.79 – 3.69 (m, 4H), 2.73 (t, *J* = 5.6 Hz, 2H), 1.49 (s, 9H) ppm. ^13^C NMR (101 MHz, CDCl_3_): *δ* = 162.1, 155.2, 141.1, 140.7, 139.9, 134.6, 130.1, 127.9, 127.7, 126.0, 116.6, 80.4, 61.0, 60.5, 50.7, 48.4, 42.4, 41.4, 40.3, 28.6, 22.1 ppm. HPLC-MS (ESI): *m/z* = 482 [M+Na]^+^, retention time: 3.87 min.

#### *tert*-Butyl 1-(2-azidoethyl)-3-((3-methylbenzyl)carbamoyl)-1,4,6,7-tetrahydro-5*H*-pyrazolo[4,3-*c*]pyridine-5-carboxylate (12)

Pyrazole azide **7** (1.13 g, 3.10 mmol, 1.0 equiv), 3-methylbenzylamine (467 μL, 451 mg, 3.72 mmol, 1.2 equiv), TBD (432 mg, 3.10 mmol, 1.0 equiv), dry THF (25.0 mL). Yield: 49% (672 mg, 1.53 mmol), colorless gum. ^1^H NMR (400 MHz, CDCl_3_): *δ* = 7.23 (t, *J* = 7.5 Hz, 1H), 7.20 – 7.12 (m, 3H), 7.09 (d, *J* = 7.5 Hz, 1H), 4.71 (s, 2H), 4.56 (d, *J* = 5.9 Hz, 2H), 4.11 (t, *J* = 5.3 Hz, 2H), 3.78 – 3.66 (m, 4H), 2.72 (t, *J* = 5.8 Hz, 2H), 2.34 (s, 3H), 1.48 (s, 9H) ppm. ^13^C NMR (101 MHz, CDCl_3_): *δ* = 161.9, 155.3, 141.3, 140.0, 138.6, 138.4, 128.8 (2x C), 128.4, 125.1, 116.7, 80.4, 50.8, 48.4, 43.1, 41.5, 28.6, 27.1, 22.2, 21.5 ppm. HPLC-MS (ESI): *m/z* = 462 [M+Na]^+^, retention time: 3.73 min.

#### *tert*-Butyl1-(2-azidoethyl)-3-((3-cyclopropylbenzyl)carbamoyl)-1,4,6,7-tetrahydro-5*H*-pyrazolo[4,3-*c*]pyridine-5-carboxylate (13)

Pyrazole azide **7** (796 mg, 2.18 mmol, 1.0 equiv), 3-cyclopropylbenzylamine hydrochloride (**10**) (2.18 mmol, 1.0 equiv), TBD (608 mg, 4.37 mmol, 2.0 equiv), dry THF (20.0 mL). Yield: 38% (384 mg, 0.825 mmol), colorless gum. ^1^H NMR (400 MHz, CDCl_3_): *δ* = 7.23 (t, *J* = 7.6 Hz, 1H), 7.16 – 7.05 (m, 3H), 6.97 (d, *J* = 7.7 Hz, 1H), 4.71 (s, 1H), 4.55 (d, *J* = 5.8 Hz, 2H), 4.10 (t, *J* = 5.6 Hz, 2H), 3.77 – 3.67 (m, 4H), 2.71 (t, *J* = 5.7 Hz, 2H), 1.89 (tt, *J* = 8.4, 5.1 Hz, 1H), 1.48 (s, 9H), 1.00 – 0.90 (m, 2H), 0.73 – 0.66 (m, 2H) ppm. ^13^C NMR (101 MHz, CDCl_3_): *δ* = 162.1, 155.3, 144.7, 141.4, 139.9, 138.4, 128.8, 125.7, 125.2, 124.7, 116.6, 80.4, 61.1, 50.8, 48.4, 43.2, 41.5, 28.7, 22.1, 15.5, 9.5 ppm. HPLC-MS (ESI): *m/z* = 488 [M+Na]^+^, retention time: 3.86 min.

### General procedure for the synthesis of target compounds 1-3

Boc-deprotection: Pyrazole azide precursor **11**, **12** or **13** (1.0 equiv) was dissolved in dry 1,4-dioxane under an argon atmosphere. After the addition of HCl (4.0 M in 1,4-dioxane, 14.0 equiv), the resulting reaction mixture was stirred for 4 h at rt. Afterwards, the solvent was removed under reduced pressure and the off-white residual solid was dried in vacuo. Reductive amination: The deprotected crude material was suspended in a mixture of dry DCE and DMF under argon and TEA (1.0 equiv) was added. After stirring the reaction for 30 min at rt, 6-methoxyquinoline-4-carbaldehyde (1.0 or 1.1 equiv), AcOH (1.0 equiv) as well as NaBH(OAc)_3_ (2.0 equiv) were added and the resulting mixture was stirred overnight. The following day, the reaction was quenched with a saturated aqueous solution of NaHCO_3_ (25 mL) and the aqueous phase was extracted with EtOAc (3 x 35 mL). Afterwards, the combined organic layers were washed with brine (25 mL), dried over Na_2_SO_4_, filtered and concentrated under reduced pressure. The obtained crude tertiary amine was used for the next step without further purification. Azide reduction: An aqueous slurry suspension of Raney-nickel (1 x tip of a spatula) was washed under argon with MeOH (3 x 5.0 mL). Afterwards, a solution of the crude azide in MeOH was added under argon. After the addition of 50% hydrazine monohydrate solution (3.2 equiv), the reaction mixture was heated until reflux and stirred for 1 h. Afterwards, Raney-nickel was filtered off via Celite^®^, washed with MeOH (25 mL) and the solvent was removed under reduced pressure. The residue was purified by gradient flash column chromatography (CH_2_Cl_2_/NH_3_ (7.0 M in MeOH): 1.5, 2.5, 3.5, 4, 5, 7% NH_3_ (7.0 M in MeOH)) and thereby the desired target compound **1**, **2** or **3** was obtained, respectively.

#### 1-(2-Aminoethyl)-*N*-(3-chlorobenzyl)-5-((6-methoxyquinolin-4-yl)methyl)-4,5,6,7-tetrahydro-1*H*-pyrazolo[4,3-*c*]pyridine-3-carboxamide (1)

Boc-deprotection: Pyrazole azide **11** (1.20 g, 2.61 mmol, 1.0 equiv), HCl (4.0 M in 1,4-dioxane, 9.13 mL, 36.5 mmol, 14.0 equiv), dry 1,4-dioxane (12.0 mL). Reductive amination: TEA (362 μL, 2.61 mmol, 1.0 equiv), 6-methoxyquinoline-4-carbaldehyde (488 mg, 2.61 mmol, 1.0 equiv), AcOH (149 μL, 2.61 mmol, 1.0 equiv), NaBH(OAc)_3_ (1.11 g, 5.22 mmol, 2.0 equiv), dry DCE/DMF (17.0/3.00 mL). Azide reduction: Hydrazine monohydrate solution (50% in H_2_O, 809 μL, 8.35 mmol, 3.2 equiv), Raney-Ni (1 x tip of a spatula), MeOH (20.0 mL). Yield: 59% (772 mg, 1.53 mmol), white foam. ^1^H NMR (400 MHz, CDCl_3_): *δ* = 8.72 (d, *J* = 4.4 Hz, 1H), 8.02 (d, *J* = 9.2 Hz, 1H), 7.49 (d, *J* = 2.8 Hz, 1H), 7.42 (d, *J* = 4.4 Hz, 1H), 7.36 (dd, *J* = 9.2, 2.8 Hz, 1H), 7.33 (s, 1H), 7.28 – 7.15 (m, 4H), 4.55 (d, *J* = 6.2 Hz, 2H), 4.12 (s, 2H), 4.02 (t, *J* = 5.8 Hz, 2H), 3.98 (s, 2H), 3.90 (s, 3H), 3.14 (t, *J* = 5.8 Hz, 2H), 2.81 (t, *J* = 5.6 Hz, 2H), 2.71 (t, *J* = 5.6 Hz, 2H) ppm. ^13^C NMR (101 MHz, CDCl_3_): *δ* = 162.8, 157.7, 147.9, 144.7, 142.6, 140.9, 140.7, 139.2, 134.7, 131.6, 130.1, 128.7, 128.0, 127.7, 126.1, 121.6 (2x C), 117.4, 102.5, 58.9, 55.7, 52.3, 50.7, 49.3, 42.4, 41.9, 22.7 ppm. HPLC-MS (ESI): *m/z* = 505 [M+H]^+^, retention time: 2.67 min. HRMS (ESI): *m/z* calcd. for [C_27_H_29_N_6_O_2_ClNa]^+^: 527.1938; found: 527.1926.

#### 1-(2-Aminoethyl)-5-((8-methoxyquinolin-4-yl)methyl)-*N*-(3-methylbenzyl)-4,5,6,7-tetrahydro-1*H*- pyrazolo[4,3-*c*]pyridine-3-carboxamide (2)

Boc-deprotection: Pyrazole azide **12** (213 mg, 0.485 mmol, 1.0 equiv), HCl (4.0 M in 1,4-dioxane, 1.70 mL, 6.80 mmol, 14.0 equiv), dry 1,4-dioxane (7.0 mL). Reductive amination: TEA (67.2 μL, 0.485 mmol, 1.0 equiv), 6-methoxyquinoline-4-carbaldehyde (99.9 mg, 0.543 mmol, 1.0 equiv), AcOH (27.7 μL, 0.485 mmol, 1.0 equiv), NaBH(OAc)_3_ (206 mg, 0.970 mmol, 2.0 equiv), dry DCE/DMF (7.00/1.00 mL). Azide reduction: Hydrazine monohydrate solution (50% in H_2_O, 151 μL, 1.55 mmol, 3.2 equiv), Raney-Ni (1 x tip of a spatula), MeOH (9.00 mL). Yield: 65% (153 mg, 0.316 mmol), white foam. ^1^H NMR (400 MHz, CDCl_3_): *δ* = 8.69 (d, *J* = 4.4 Hz, 1H), 8.39 (t, *J* = 6.4 Hz, 1H), 7.94 (d, *J* = 9.2 Hz, 1H), 7.59 (d, *J* = 2.8 Hz, 1H), 7.49 (d, *J* = 4.3 Hz, 1H), 7.40 (dd, *J* = 9.2, 2.8 Hz, 1H), 7.16 (t, *J* = 7.5 Hz, 1H), 7.08 – 6.98 (m, 3H), 4.30 (d, *J* = 6.3 Hz, 2H), 4.14 (s, 2H), 3.98 (t, *J* = 6.3 Hz, 2H), 3.88 (s, 3H), 3.63 (s, 2H), 2.94 – 2.84 (m, 4H), 2.79 (d, *J* = 5.4 Hz, 2H), 2.25 (s, 3H) ppm. ^13^C NMR (101 MHz, DMSO-d_6_): *δ* = 162.0, 156.9, 147.6, 143.9, 142.7, 140.0, 139.8, 138.9, 137.1, 130.9, 128.0, 127.9, 127.2, 124.4, 121.3, 121.1, 115.6, 103.0, 57.8, 55.3, 51.9, 49.6, 49.4, 48.6, 41.7, 41.6, 21.7, 21.0 ppm. HPLC-MS (ESI): *m/z* = 485 [M+H]^+^, 507 [M+Na]^+^, retention time: 2.32 min. HRMS (ESI): *m/z* calcd. for [C_28_H_32_N_6_O_2_Na]^+^: 507.2484; found: 507.2463.

#### 1-(2-Aminoethyl)-*N*-(3-cyclopropylbenzyl)-5-((6-methoxyquinolin-4-yl)methyl)-4,5,6,7-tetrahydro-1*H*-pyrazolo[4,3-*c*]pyridine-3-carboxamide (3)

Boc-deprotection: Pyrazole azide **13** (170 mg, 0.365 mmol, 1.0 equiv), HCl (4.0 M in 1,4-dioxane, 1.28 mL, 5.11 mmol, 14.0 equiv), dry 1,4-dioxane (7.00 mL). Reductive amination: TEA (50.6 μL, 0.365 mmol, 1.0 equiv), 6-methoxyquinoline-4-carbaldehyde (71.7 mg, 0.383 mmol, 1.1 equiv), AcOH (20.9 μL, 0.365 mmol, 1.0 equiv), NaBH(OAc)_3_ (155 mg, 0.730 mmol, 2.0 equiv), dry DCE/DMF (6.00/1.00 mL). Azide reduction: Hydrazine monohydrate solution (50% in H_2_O, 113 μL, 1.17 mmol, 3.2 equiv), Raney-Ni (1 x tip of a spatula), MeOH (9.00 mL). Yield: 61% (114 mg, 0.224 mmol), white foam. ^1^H NMR (400 MHz, CDCl_3_): *δ* = 8.69 (d, *J* = 4.3 Hz, 1H), 8.38 (t, *J* = 6.3 Hz, 1H), 7.94 (d, *J* = 9.2 Hz, 1H), 7.59 (d, *J* = 2.7 Hz, 1H), 7.49 (d, *J* = 4.4 Hz, 1H), 7.40 (dd, *J* = 9.2, 2.8 Hz, 1H), 7.14 (t, *J* = 7.6 Hz, 1H), 7.04 – 6.94 (m, 2H), 6.88 (d, *J* = 7.7 Hz, 1H), 4.29 (d, *J* = 6.3 Hz, 2H), 4.14 (s, 2H), 3.98 (t, *J* = 6.2 Hz, 2H), 3.88 (s, 3H), 3.64 (s, 2H), 2.95 – 2.83 (m, 4H), 2.83 – 2.74 (m, 2H), 1.85 (tt, *J* = 8.4, 5.9 Hz, 1H), 0.95 – 0.81 (m, 2H), 0.65 – 0.53 (m, 2H) ppm. ^13^C NMR (101 MHz, DMSO-d_6_): *δ* = 162.0, 156.9, 147.6, 143.9, 143.4, 142.7, 140.0, 139.8, 138.9, 130.9, 128.1, 124.7, 124.3, 123.4 (2x C), 121.3, 121.1, 115.6, 103.0, 57.7, 55.3, 51.9, 49.6, 49.4, 41.7, 41.7, 21.7, 14.9, 9.2 ppm. HPLC-MS (ESI): *m/z* = 511 [M+H]^+^, 533 [M+Na]^+^, retention time: 2.49 min. HRMS (ESI): *m/z* calcd. for [C_30_H_34_N_6_O_2_Na]^+^: 533.2641; found: 507.2657.

### Synthetic procedures and spectroscopic data for PEX14 inhibitors 4-5

**Scheme S2.**
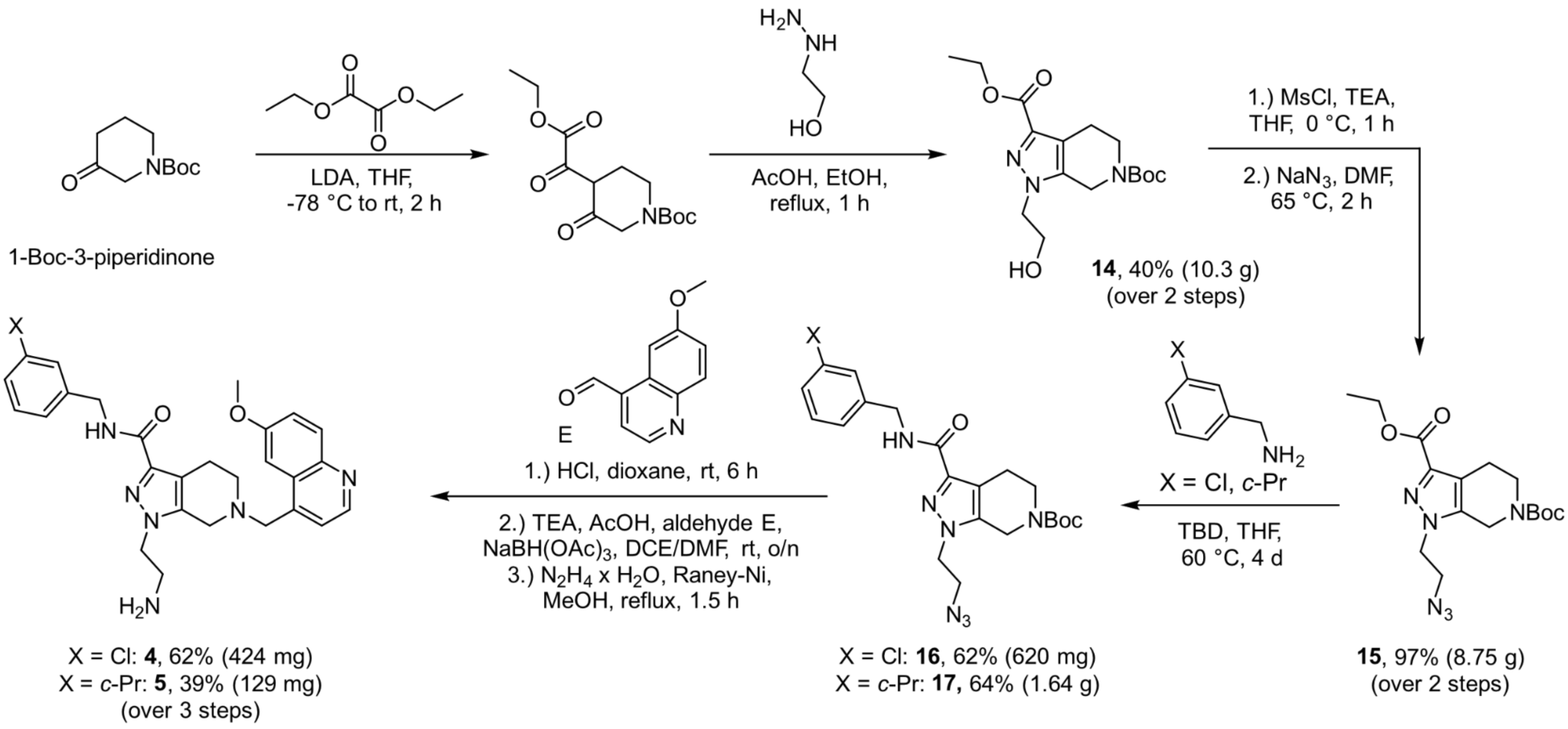
Synthetic route for the preparation of PEX14 inhibitors **4-5**.

#### 6-(*tert*-Butyl) 3-ethyl 1-(2-hydroxyethyl)-1,4,5,7-tetrahydro-6*H*-pyrazolo[3,4-*c*]pyridine-3,6-dicarboxylate (14)

A solution of lithium diisopropylamide (LDA) (2.0 M in THF, 41.4 mL, 82.8 mmol, 1.1 equiv) in dry THF (100 mL) was cooled to -78 °C. To this mixture, a solution of 1-Boc-3-piperidone (15.0 g, 75.3 mmol, 1.0 equiv) in dry THF (50.0 mL) was added dropwise under an argon atmosphere. After 30 min of continuous stirring at -78 °C, a solution of diethyl oxalate (10.2 mL, 75.3 mmol, 1.0 equiv) in dry THF (50.0 mL) was slowly added. The resulting reaction mixture was then warmed to rt and stirred for an additional 30 min. Subsequently, 2-hydroxyethylhydrazine (5.10 mL, 75.3 mmol, 1.0 equiv), acetic acid (25.0 mL) and EtOH (40.0 mL) were added. After refluxing for 1 h, the reaction was cooled to rt and concentrated in vacuo. The remaining residue was partitioned between EtOAc (150 mL) and a saturated NaHCO_3_ solution (100 mL). The aqueous phase was extracted with EtOAc (3 × 100 mL) and the combined organic extracts were washed with water (75.0 mL) and brine (75.0 mL). After drying over Na_2_SO_4_, the extract was filtered and concentrated in vacuo. The obtained oily residue was then purified via gradient column chromatography (*c*-hexane/EtOAc: 10, 25, 35, 50, 75, 100% EtOAc to yield precursor **14** as an off-white solid (10.3 g, 30.3 mmol, 40%). ^1^H NMR (400 MHz, CDCl_3_): *δ* = 4.57 (s, 2H), 4.37 (q, *J* = 7.2 Hz, 2H), 4.22 (t, *J* = 5.0 Hz, 2H), 4.04 (t, *J* = 5.1 Hz, 2H), 3.62 (t, *J* = 5.8 Hz, 2H), 3.58 (br s, 1H), 2.83 (t, J = 5.8 Hz, 2H), 1.47 (s, 9H), 1.38 (t, J = 7.1 Hz, 3H) ppm. ^13^C NMR (101 MHz, CDCl_3_): *δ* = 162.3, 154.9, 139.4, 138.2, 118.3, 80.6, 61.3, 60.8, 52.0, 40.4, 28.3, 21.9, 14.3 ppm. HPLC-MS (ESI): *m/z* = 362 [M+Na]^+^; retention time: 3.26 min.

#### 6-(*tert*-Butyl) 3-ethyl 1-(2-azidoethyl)-1,4,6,7-tetrahydro-6*H*-pyrazolo[3,4-*c*]pyridine-3,6-dicarboxylate (15)

Pyrazole **14** (8.40 g, 24.8 mmol, 1.0 eq) was dissolved in dry THF (50.0 mL) under an argon atmosphere and cooled to 0 °C. After the addition of triethylamine (6.86 mL, 49.5 mmol, 2.0 equiv) and methanesulfonyl chloride (2.49 mL, 32.2 mmol, 1.3 equiv), the resulting mixture was stirred for 1 h at 0 °C. Afterwards, the reaction mixture was filtered through Celite^®^ and the solvent was removed under reduced pressure. For the second step, the crude mesylate was redissolved in dry DMF (25.0 mL) and sodium azide (4.83 g, 75.3 mmol, 3.0 equiv) was added. The mixture was then heated to 70 °C for 2 h, after which it was quenched by addition of a saturated NaHCO_3_ solution (50.0 mL). The aqueous phase was then extracted with EtOAc (3 × 70.0 mL) and the combined organic phases were washed with water (50.0 mL) and brine (50.0 mL), dried over Na_2_SO_4_, filtered and concentrated under reduced pressure. The crude product was purified via gradient column chromatography (*c*-hexane/EtOAc: 25, 35, 50, 75, 100% EtOAc), which yielded the desired product **15** as an off-white solid (8.75 g, 24.0 mmol, 97%). ^1^H NMR (400 MHz, CDCl_3_): *δ* = 4.56 (s, 2H), 4.41 (q, *J* = 7.1 Hz, 2H), 4.17 (t, J = 5.5 Hz, 2H), 3.79 (t, J = 5.5 Hz, 2H), 3.62 (t, J = 5.7 Hz, 2H), 2.83 (t, J = 5.7 Hz, 2H), 1.48 (s, 9H), 1.38 (t, J = 7.1, 0.7 Hz, 3H) ppm. ^13^C NMR (101 MHz, CDCl_3_): *δ* = 162.5, 155.0, 140.5, 138.3, 118.7, 80.8, 61.0, 51.2, 49.3, 40.5, 28.5, 27.1, 22.1, 14.6 ppm. HPLC-MS (ESI): *m/z* = 387 [M+Na]^+^, retention time: 3.69 min.

#### General procedure for the synthesis of precursors 16 and 17

A solution of pyrazole azide **15** (1.0 equiv), the respective benzylamine (1.2 equiv) and TBD (1.0 or 2.0 equiv) in dry THF was stirred at 60 °C for 4 d. The mixture was concentrated *in vacuo* and the residue was purified by gradient flash column chromatography (*c*-hexane/EtOAc: 20, 35, 50, 70, 90, 100% EtOAc).

#### *tert*-Butyl 1-(2-azidoethyl)-3-((3-chlorobenzyl)carbamoyl)-1,4,5,7-tetrahydro-6*H*-pyrazolo[3,4-*c*]pyridine-6-carboxylate (16)

Pyrazole azide **15** (2.00 g, 5.49 mmol, 1.0 equiv), 3-chlorobenzylamine (805 μL, 933 mg, 6.59 mmol, 1.2 equiv), TBD (764 mg, 5.49 mmol, 1.0 equiv), dry THF (25.0 mL). Yield: 62% (1.57 g, 3.41 mmol), white solid. ^1^H NMR (400 MHz, CDCl_3_): *δ* = 7.34 – 7.32 (m, 1H), 7.27 – 7.20 (m, 4H), 4.59 – 4.50 (m, 4H), 4.09 (t, *J* = 5.2 Hz, 2H), 3.76 (t, *J* = 5.7 Hz, 2H), 3.62 (t, *J* = 5.5 Hz, 2H), 2.92 (t, *J* = 5.6 Hz, 2H), 1.48 (s, 9H) ppm. ^13^C NMR (101 MHz, CDCl_3_): *δ* = 162.3, 142.1, 140.7, 138.5, 134.7, 130.2, 130.1, 128.0, 127.7, 126.1, 117.5, 80.8, 50.7, 49.0, 43.3, 42.5, 28.6, 23.3, 21.9 ppm. HPLC-MS (ESI): *m/z* = 482 [M+Na]^+^, retention time: 3.80 min.

#### *tert*-Butyl 1-(2-azidoethyl)-3-((3-cyclopropylbenzyl)carbamoyl)-1,4,5,7-tetrahydro-6*H*-pyrazolo[3,4-*c*]pyridine-6-carboxylate (17)

Pyrazole azide **15** (2.00 g, 5.49 mmol, 1.0 equiv), crude 3-cyclopropylbenzylamine hydrochloride (**10**) (6.59 mmol, 1.2 equiv), TBD (1.53 g, 10.98 mmol, 2.0 equiv), dry THF (25.0 mL). Yield: 64% (1.64 g, 3.52 mmol), colorless gum. ^1^H NMR (400 MHz, CDCl_3_): *δ* = 7.24 – 7.20 (t, *J* = 7.6 Hz, 1H), 7.14 – 7.11 (m, 1H), 7.07 – 7.06 (m, 1H), 6.97 – 6.95 (m, 1H), 4.55 – 4.54 (m, 4H), 4.07 (t, *J* = 5.5 Hz, 2H), 3.74 (t, *J* = 5.6 Hz, 2H), 3.62 (t, *J* = 5.4 Hz, 2H), 2.93 (t, *J* = 5.9 Hz, 2H), 1.91 – 1.85 (tt, *J* = 8.4, 5.1 Hz, 1H), 1.49 (s, 9H), 1.01 – 0.88 (m, 2H), 0.69 (dt, *J* = 6.7, 4.6 Hz, 2H) ppm. ^13^C NMR (101 MHz, CDCl_3_): *δ* = 162.2, 155.2, 144.7, 142.4, 138.5, 138.4, 128.8, 125.7, 125.2, 124.7, 117.2, 80.7, 50.7, 48.9, 43.1, 40.5, 28.6, 21.9, 15.5, 9.4 ppm. HPLC-MS (ESI): *m/z* = 359 [M-Boc]^+^, retention time: 3.26 min.

#### General procedure for the synthesis of target compounds 4 and 5

Boc-deprotection: Pyrazole azide precursor **16** or **17** (1.0 equiv) was dissolved in dry 1,4-dioxane under an argon atmosphere. After the addition of HCl (4.0 M in 1,4-dioxane, 14.0 equiv), the resulting reaction mixture was stirred for 6 h at rt. Afterwards, the solvent was removed under reduced pressure and the off-white residual solid was dried in vacuo. Reductive amination: The deprotected crude material was suspended in a mixture of dry DCE and DMF under argon and TEA (1.0 equiv) was added. After stirring the reaction for 30 min at rt, 6-methoxyquinoline-4-carbaldehyde (1.1 equiv), AcOH (1.0 equiv) as well as NaBH(OAc)_3_ (2.0 equiv) were added and the resulting mixture was stirred overnight. The following day, the reaction was quenched with a saturated aqueous solution of NaHCO_3_ (25 mL) and the aqueous phase was extracted with EtOAc (3 x 35 mL). Afterwards, the combined organic layers werewashed with brine (25 mL), dried over Na_2_SO_4_, filtered and concentrated under reduced pressure. The obtained crude tertiary amine was used for the next step without further purification. Azide reduction: An aqueous slurry suspension of Raney-nickel (1 x tip of a spatula) was washed under argon with MeOH (3 x 5.0 mL). Afterwards, a solution of the crude azide in MeOH was added under argon. After the addition of 50% hydrazine monohydrate solution (3.2 equiv), the reaction mixture was heated until reflux and stirred for 1 h. Afterwards, Raney-nickel was filtered off via Celite^®^, washed with MeOH (25 mL) and the solvent was removed under reduced pressure. The residue was purified by gradient flash column chromatography (CH_2_Cl_2_/NH_3_ (7.0 M in MeOH): 1.5, 2, 2.5, 3, 4, 5% NH_3_ (7.0 M in MeOH)) and thereby the desired target compound **4** or **5** was obtained, respectively.

#### 1-(2-Aminoethyl)-*N*-(3-chlorobenzyl)-6-((6-methoxyquinolin-4-yl)methyl)-4,5,6,7-tetrahydro-1*H*-pyrazolo[3,4-*c*]pyridine-3-carboxamide (4)

Boc-deprotection: Pyrazole azide **16** (620 mg, 1.35 mmol, 1.0 equiv), HCl (4.0 M in 1,4-dioxane, 4.72 mL, 18.9 mmol, 14.0 equiv), dry 1,4-dioxane (10.0 mL). Reductive amination: TEA (187 μL, 1.35 mmol, 1.0 equiv), 6-methoxyquinoline-4-carbaldehyde (265 mg, 1.42 mmol, 1.1 equiv), AcOH (77.1 μL, 1.35 mmol, 1.0 equiv), NaBH(OAc)_3_ (571 mg, 2.70 mmol, 2.0 equiv), dry DCE/DMF (12.0/2.00 mL). Azide reduction: Hydrazine monohydrate solution (50% in H_2_O, 423 μL, 4.31 mmol, 3.2 equiv), Raney-Ni (1 x tip of a spatula), MeOH (15.0 mL). Yield: 62% (424 mg, 0.840 mmol), white foam. ^1^H NMR (400 MHz, DMSO-d_6_): *δ* = 8.70 (d, *J* = 4.3 Hz, 1H), 8.56 (t, *J* = 6.4 Hz, 1H), 7.94 (d, *J* = 9.2 Hz, 1H), 7.55 (d, *J* = 2.8 Hz, 1H), 7.50 (d, *J* = 4.4 Hz, 1H), 7.40 (dd, *J* = 9.2, 2.8 Hz, 1H), 7.38 – 7.31 (m, 2H), 7.30 – 7.22 (m, 2H), 4.38 (d, *J* = 6.3 Hz, 2H), 4.15 (s, 2H), 3.98 – 3.83 (m, 5H), 3.71 (s, 2H), 2.87 (t, *J* = 6.2 Hz, 2H), 2.83 – 2.74 (m, 4H), 2.55 – 2.46 ppm. ^13^C NMR (101 MHz, DMSO-d_6_): *δ* = 162.3, 156.9, 147.6, 143.9, 142.7, 142.4, 140.9, 138.8, 132.8, 130.9, 130.1, 128.0, 127.1, 126.5, 126.0, 121.2, 115.0, 102.8, 57.3, 55.4, 52.1, 50.1, 48.3, 41.7, 41.2, 21.5 ppm. HPLC-MS (ESI): *m/z* = 505 [M+H]^+^, retention time: 2.61 min. HRMS (ESI): *m/z* calcd. for [C_27_H_29_N_6_O_2_ClNa]^+^: 527.1938; found: 527.1967.

#### 1-(2-Aminoethyl)-*N*-(3-cyclopropylbenzyl)-6-((6-methoxyquinolin-4-yl)methyl)-4,5,6,7-tetrahydro-1*H*-pyrazolo[3,4-*c*]pyridine-3-carboxamide (5)

Boc-deprotection: Pyrazole azide **17** (305 mg, 0.656 mmol, 1.0 equiv), HCl (4.0 M in 1,4-dioxane, 2.30 mL, 9.18 mmol, 14.0 equiv), dry 1,4-dioxane (6.00 mL). Reductive amination: TEA (90.9 μL, 0.656 mmol, 1.0 equiv), 6-methoxyquinoline-4-carbaldehyde (135 mg, 0.722 mmol, 1.1 equiv), AcOH (37.5 μL, 0.656 mmol, 1.0 equiv), NaBH(OAc)_3_ (278 mg, 1.31 mmol, 2.0 equiv), dry DCE/DMF (7.00/1.50 mL). Azide reduction: Hydrazine monohydrate solution (50% in H_2_O, 206 μL, 2.09 mmol, 3.2 equiv), Raney-Ni (1 x tip of a spatula), MeOH (10.0 mL). Yield: 39% (129 mg, 0.253 mmol), white solid. ^1^H NMR (400 MHz, DMSO-d_6_): *δ* = 8.70 (d, *J* = 4.3 Hz, 1H), 8.38 (t, *J* = 6.4 Hz, 1H), 7.94 (d, *J* = 9.2 Hz, 1H), 7.55 (d, *J* = 2.8 Hz, 1H), 7.50 (d, *J* = 4.4 Hz, 1H), 7.40 (dd, *J* = 9.2, 2.8 Hz, 1H), 7.17 (t, *J* = 7.6 Hz, 1H), 7.07 – 7.00 (m, 2H), 6.90 (d, *J* = 7.7 Hz, 1H), 4.34 (d, *J* = 6.3 Hz, 2H), 4.15 (s, 2H), 3.93 – 3.85 (m, 5H), 3.71 (s, 2H), 2.86 (t, *J* = 6.2 Hz, 2H), 2.81 – 2.71 (m, 3H), 1.87 (tt, *J* = 8.4, 5.1 Hz, 1H), 0.96 – 0.87 (m, 2H), 0.68 – 0.54 (m, 2H) ppm. ^13^C NMR (101 MHz, DMSO-d_6_): *δ* = 162.1, 156.9, 147.6, 143.9, 143.4, 142.4, 141.1, 139.9, 138.7, 130.9, 128.1, 128.0, 124.7, 124.3, 123.4, 121.2 (2x C), 114.8, 102.8, 57.3, 55.4, 52.1, 50.1, 48.3, 41.7, 41.7, 21.6, 15.0, 9.2 ppm. HPLC-MS (ESI): *m/z* = 511 [M+H]^+^, retention time: 2.77 min. HRMS (ESI): *m/z* calcd. for [C_30_H_34_N_6_O_2_Na]^+^: 533.2641; found: 533.2632.

### PEX14 protein expression and purification

N-terminal domain of *T. brucei*, *T. cruzi* and *H. sapiens* PEX14 (residues 19-84) were cloned into pET21 vector. The plasmids were transformed into E. coli BL21(DE3). The overnight culture was inoculated in autoinduction medium supplemented with 100 µg ml^-1^ kanamycin.^65^ When the cell density (OD600) reached 0.8, the temperature was lowered to 18°C and the cells were grown overnight. Cells were harvested by centrifugation and dissolved in lysis buffer (50 mM Tris pH 8.0, 300 mM NaCl, 10 mM b-mercaptoethanol, 20 mM imidazole, 10 mg/ml DNAseI, 1 mM AEBSF) and lysed by sonication. The lysates, clarified by centrifugation, were passed over a Ni-NTA agarose resin (Qiagen, Germany) pre-equilibrated with buffer A (50 mM Tris pH 8.0, 300 mM NaCl, 10 mM b -mercaptoethanol, 20 mM imidazole) and the protein of interest was eluted with the same buffer containing 250mM imidazole. Concentrated eluates were further purified on Superdex 75 Hiload 16/60 column (GE Healthcare) in phosphate buffer saline (PBS).

### AlphaScreen Assay

The assay was performed according to published protocols.^38,39^ 3 nM N-His-PEX14 from each species (*T. brucei*, *T. cruzi* and *H. sapiens*) was mixed with 10 nM biotinylated PEX5-derived peptide (ALSENWAQEFLA) in a PBS buffer supplemented with 5 mg/mL of BSA and 0.01 % (v/v) Tween-20. 5 µg/mL of streptavidin donor beads and 5 µg/mL of nickel chelate acceptor beads (PerkinElmer) were added to the mixture. Serial dilutions of the inhibitors (600-0 µM) were prepared in DMSO and added to the protein-peptide mixture keeping constant the concentration of DMSO (5%). Experiments were prepared in triplicates using a liquid handler OT-2 robot (Opentrons). The AlphaScreen assay was performed in white 384-well Optiplates (PerkinElmer, product no. 6007299). The AlphaScreen signals were detected at 520–620 nm using laser excitation at 680 nm, in an EnVision 2102 Multilabel Reader (PerkinElmer). Data were analyzed using Origin Pro 8.5. Experimental points were integrated using Hill sigmoidal fitting fixing the asymptotes at the maximal assay signal (no inhibitor added) and 0, respectively.

#### Cellular model of Trypanosoma cruzi infection

T.cruzi Tulahuen strain C2C4 containing the β-galactosidase (Lac Z) gene^34^ as cultivated using Mouse Embryo Fibroblasts (MEF-cells) as host cells, in RPMI 1640 medium supplemented with 1% L-glutamine (200mM) and 10% fetal bovine serum. Cultures were maintained at 37 °C in an atmosphere of 5% CO2. Rat skeletal myoblasts (L-6 cells)^35,36^ were cultivated in RPMI 1640 medium supplemented with 1% L-glutamine (200mM) and 10% fetal bovine serum. L-6 Cultures were maintained at 37 °C in an atmosphere of 5% CO2. Assays were performed in 96-well plates containing RPMI 1640 medium supplemented with 1% L-glutamine (200 mM), 10% fetal bovine serum, and 1×10^3^ L-6 cells/well. Plates were incubated at 37 °C under a 5% CO2 atmosphere for 24h. After 24 h, trypomastigote forms of T.cruzi Tulahuen strain C2C4 containing the β-galactosidase (LacZ) gene were added to the wells (5×10^3^ /well) for infection. Compounds were dissolved in DMSO (10 mg/ml) and 48h after infection serial drug dilutions of eleven 3-fold dilution steps covering 100 to 0.002 μg/mL were prepared in fresh 96 well plates. Benznidazole was used as a reference compound. The medium from the assay plates was removed and replaced by 100 μl of the serial drug dilution plate. After 96h the substrate CPRG/IGEPAL CA 630 (50 μl /well) was added to all wells. A color reaction developed within 2–6h and was read photometrically at 540 nm using the TECAN Sparks (Tecan). IC50 values were calculated from sigmoidal dose response curves by linear regression^37^ using Microsoft Excel. Two independent replicates of the assay were performed.

##### 1H and ^13^C NMR Spectra of Inhibitors 1-5

^1^H NMR spectrum of compound **1** recorded on a Bruker AV-HD400 spectrometer (400 MHz, CDCl_3_):

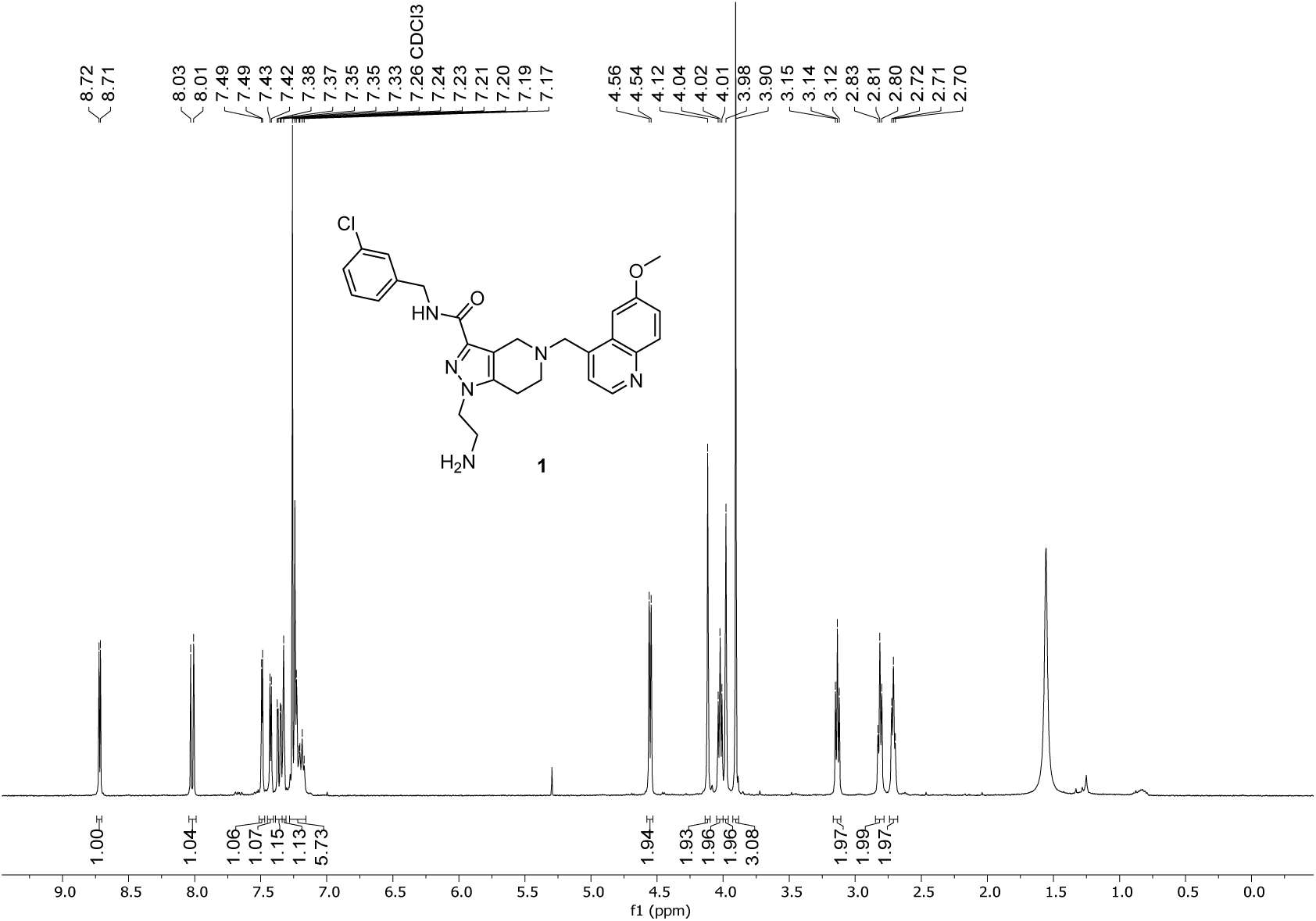

^13^C NMR spectrum of compound **1** recorded on a Bruker AV-HD400 spectrometer (101 MHz, CDCl_3_):

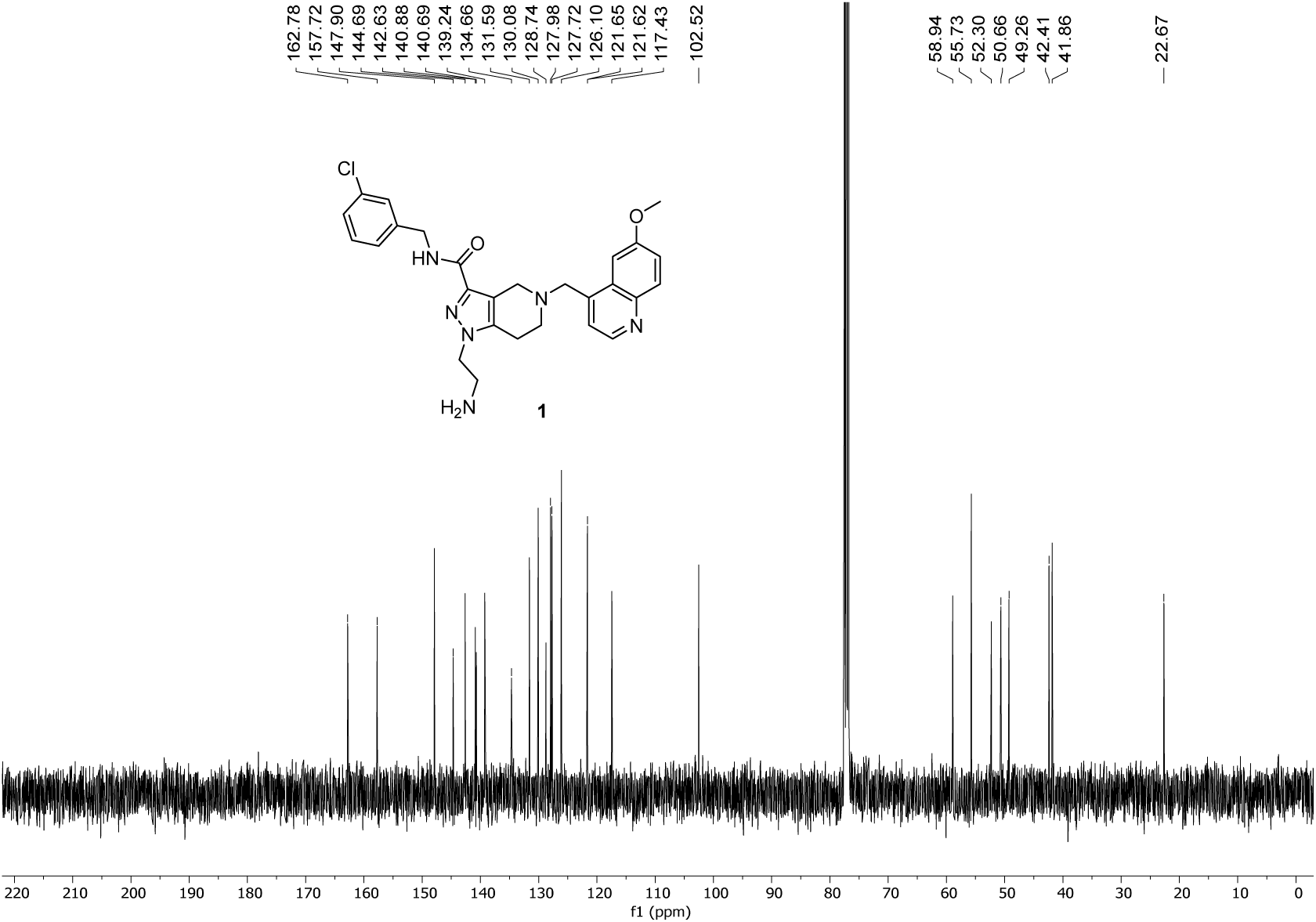

^1^H NMR spectrum of compound **2** recorded on a Bruker AV-HD400 spectrometer (400 MHz, CDCl_3_):

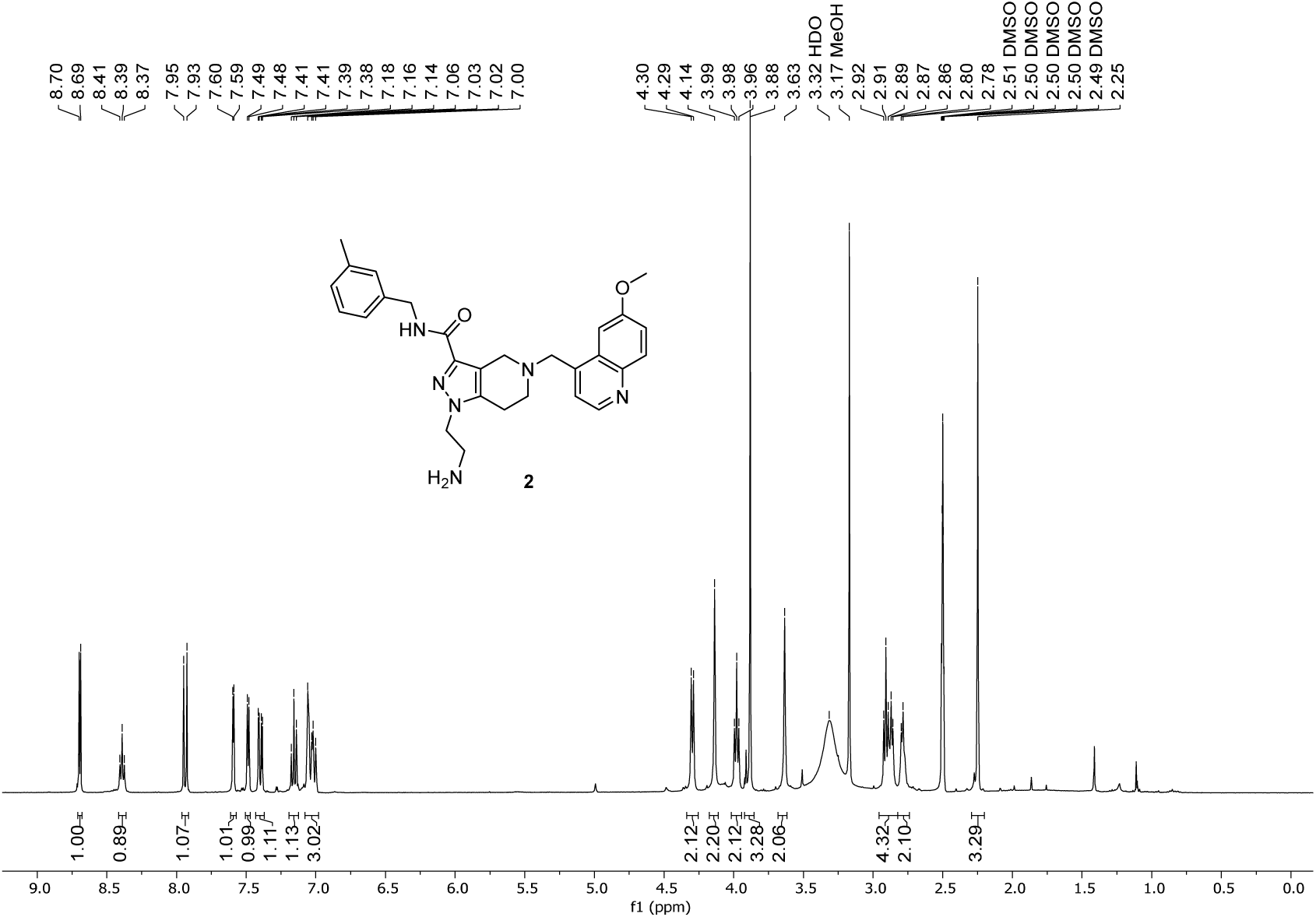

^13^C NMR spectrum of compound **2** recorded on a Bruker AV-HD400 spectrometer (101 MHz, CDCl_3_):

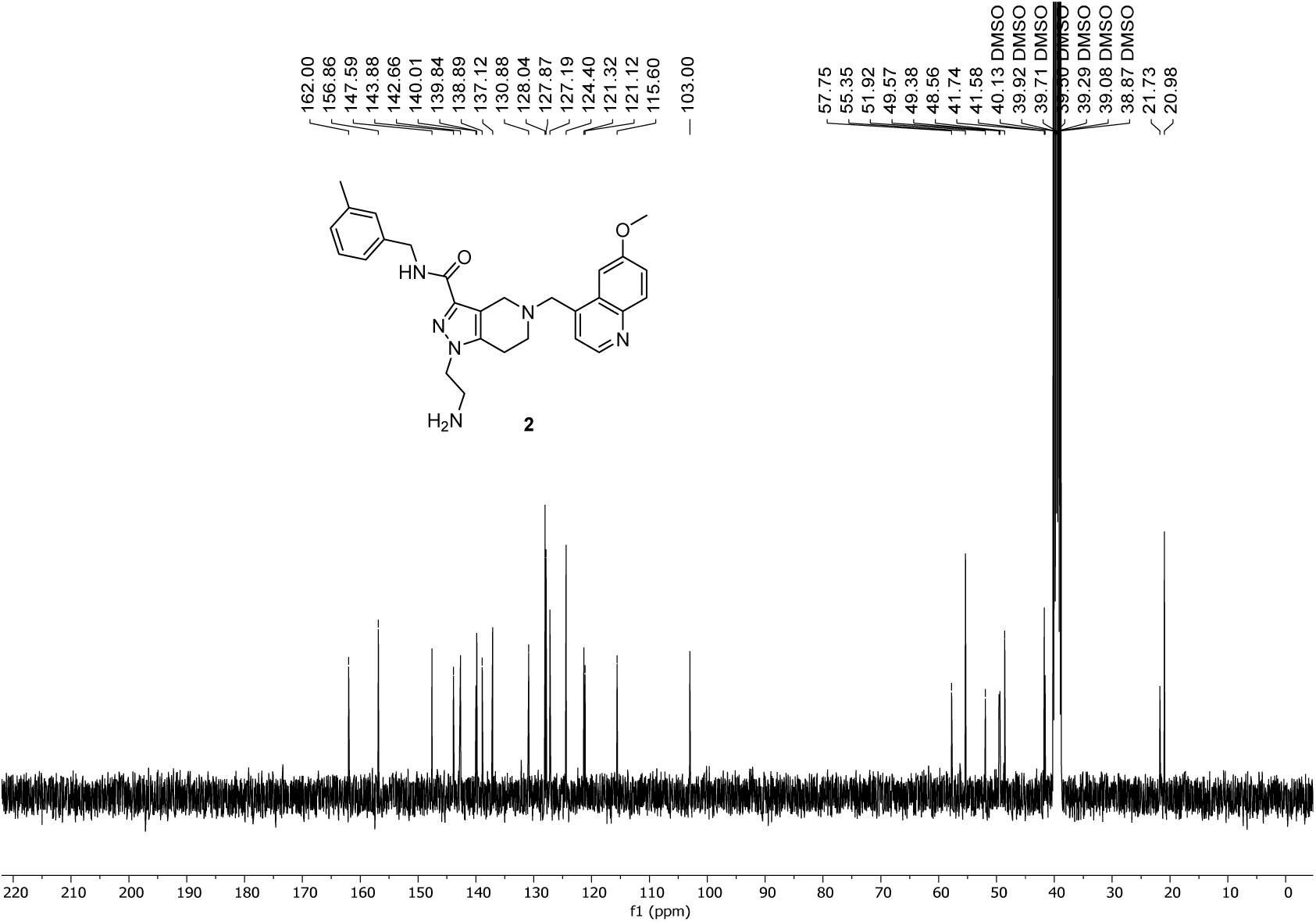

^1^H NMR spectrum of compound **3** recorded on a Bruker AV-HD400 spectrometer (400 MHz, CDCl_3_):

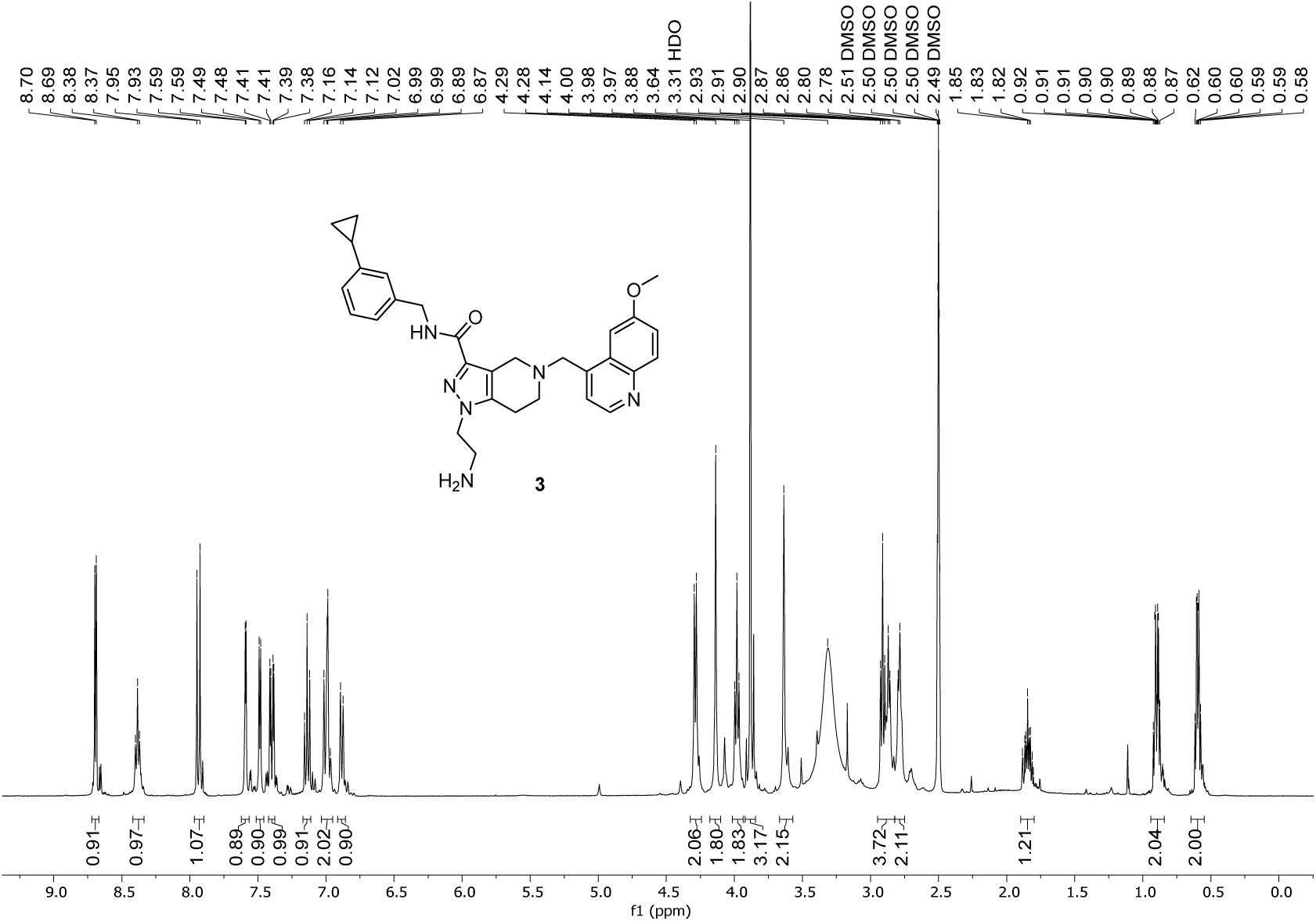

^13^C NMR spectrum of compound **3** recorded on a Bruker AV-HD400 spectrometer (101 MHz, CDCl_3_):

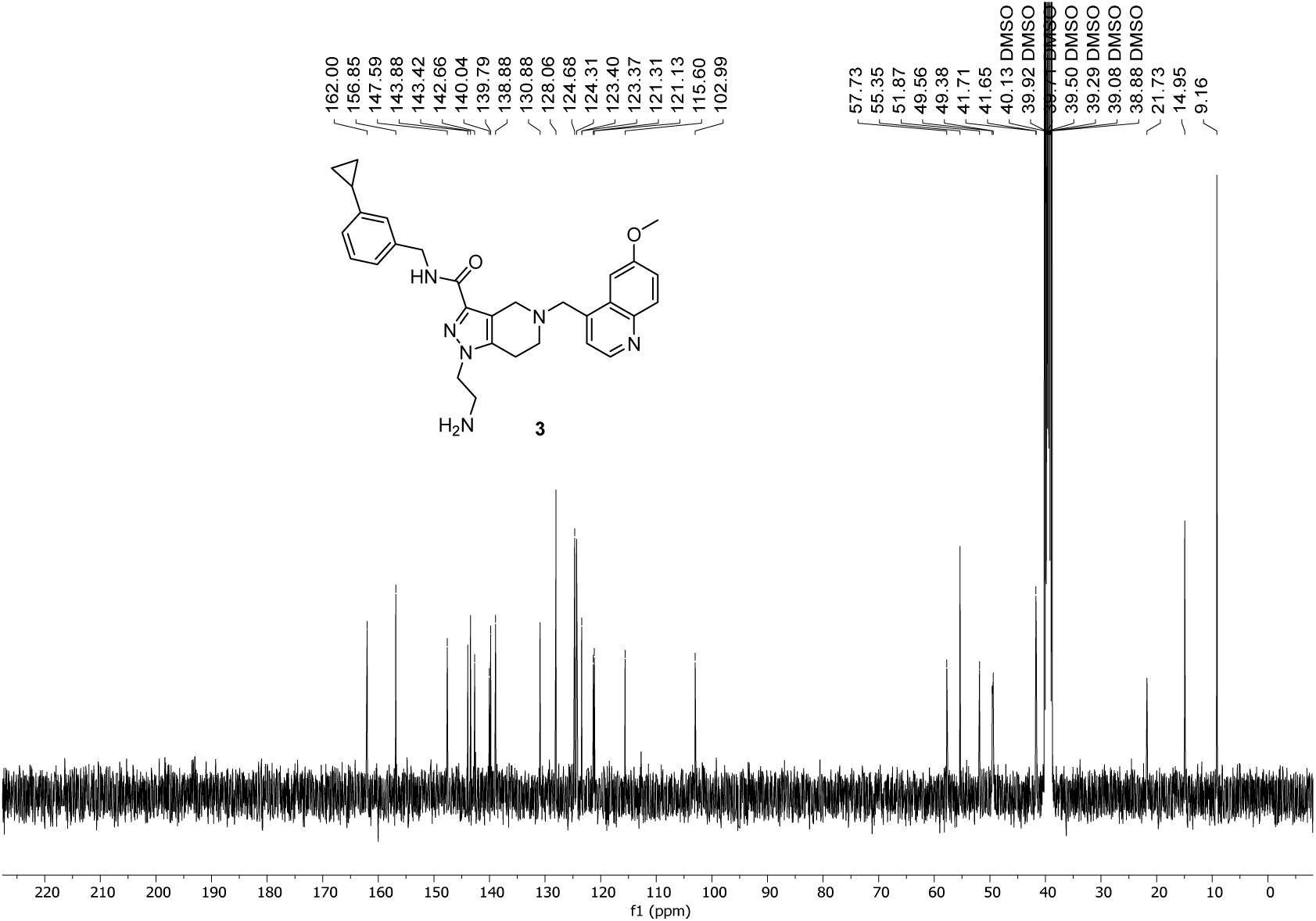

^1^H NMR spectrum of compound **4** recorded on a Bruker AV-HD400 spectrometer (400 MHz, CDCl_3_):

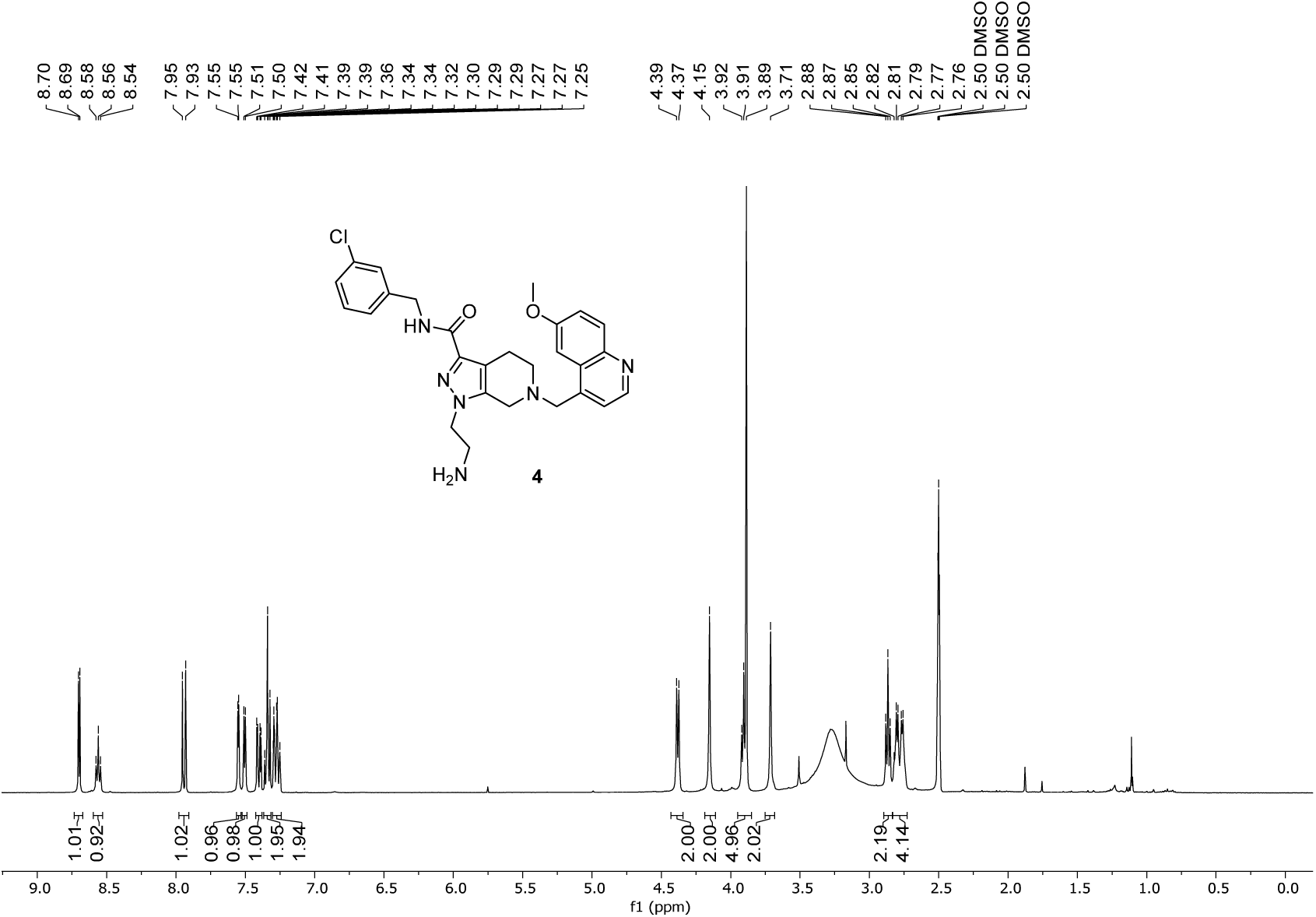

^13^C NMR spectrum of compound **4** recorded on a Bruker AV-HD400 spectrometer (101 MHz, CDCl_3_):

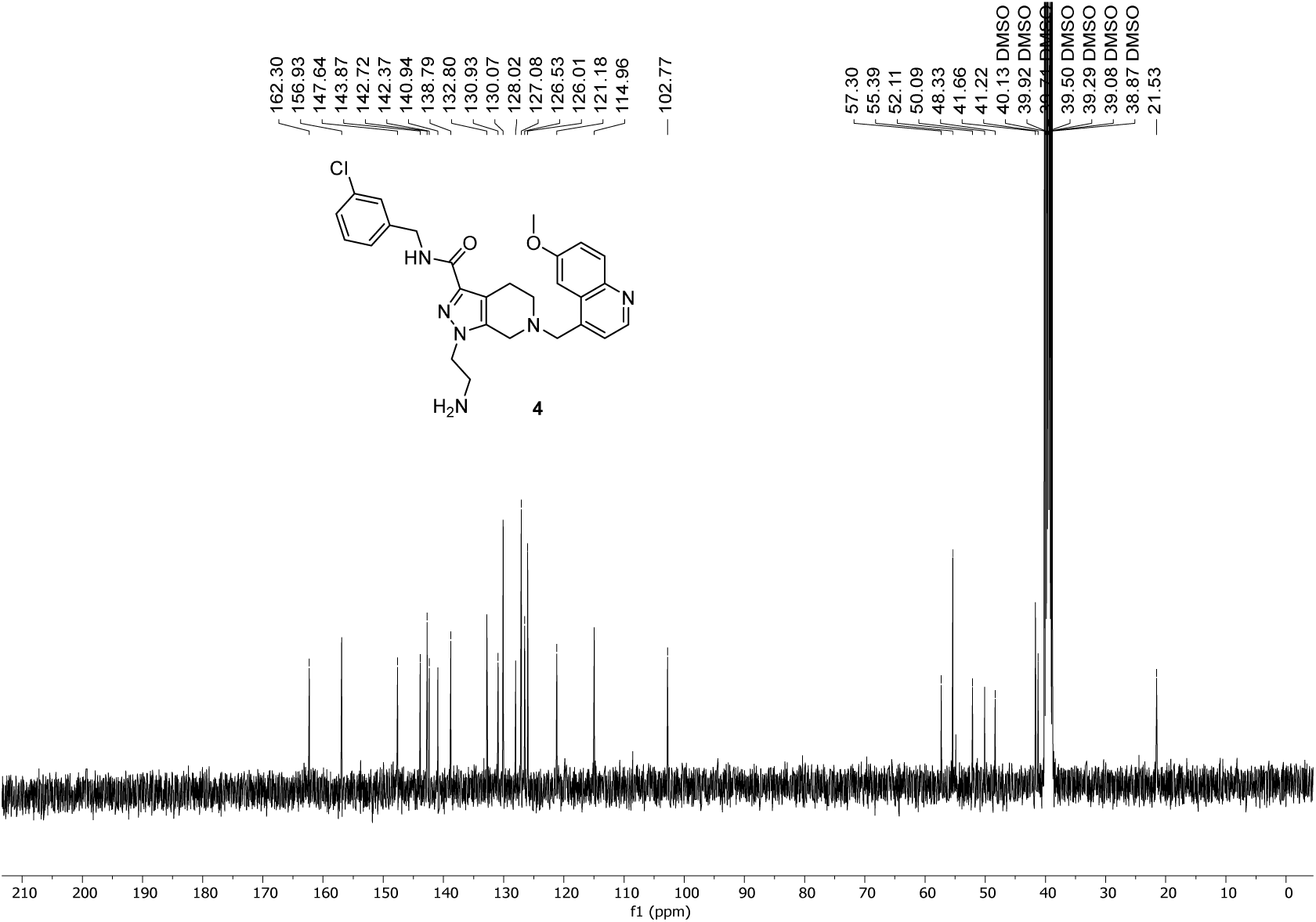

^1^H NMR spectrum of compound **5** recorded on a Bruker AV-HD400 spectrometer (400 MHz, CDCl_3_):

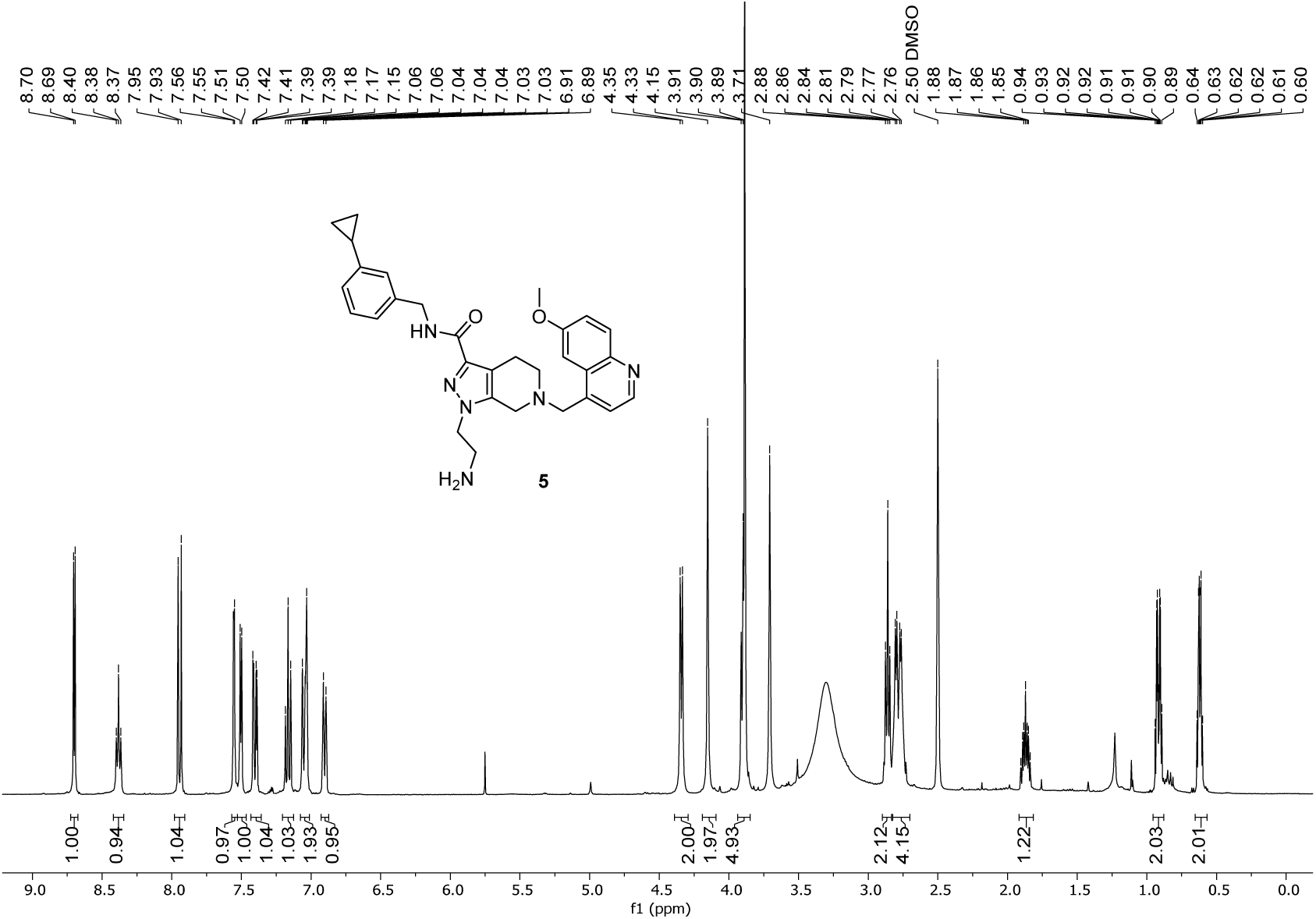

^13^C NMR spectrum of compound **5** recorded on a Bruker AV-HD400 spectrometer (101 MHz, CDCl_3_):

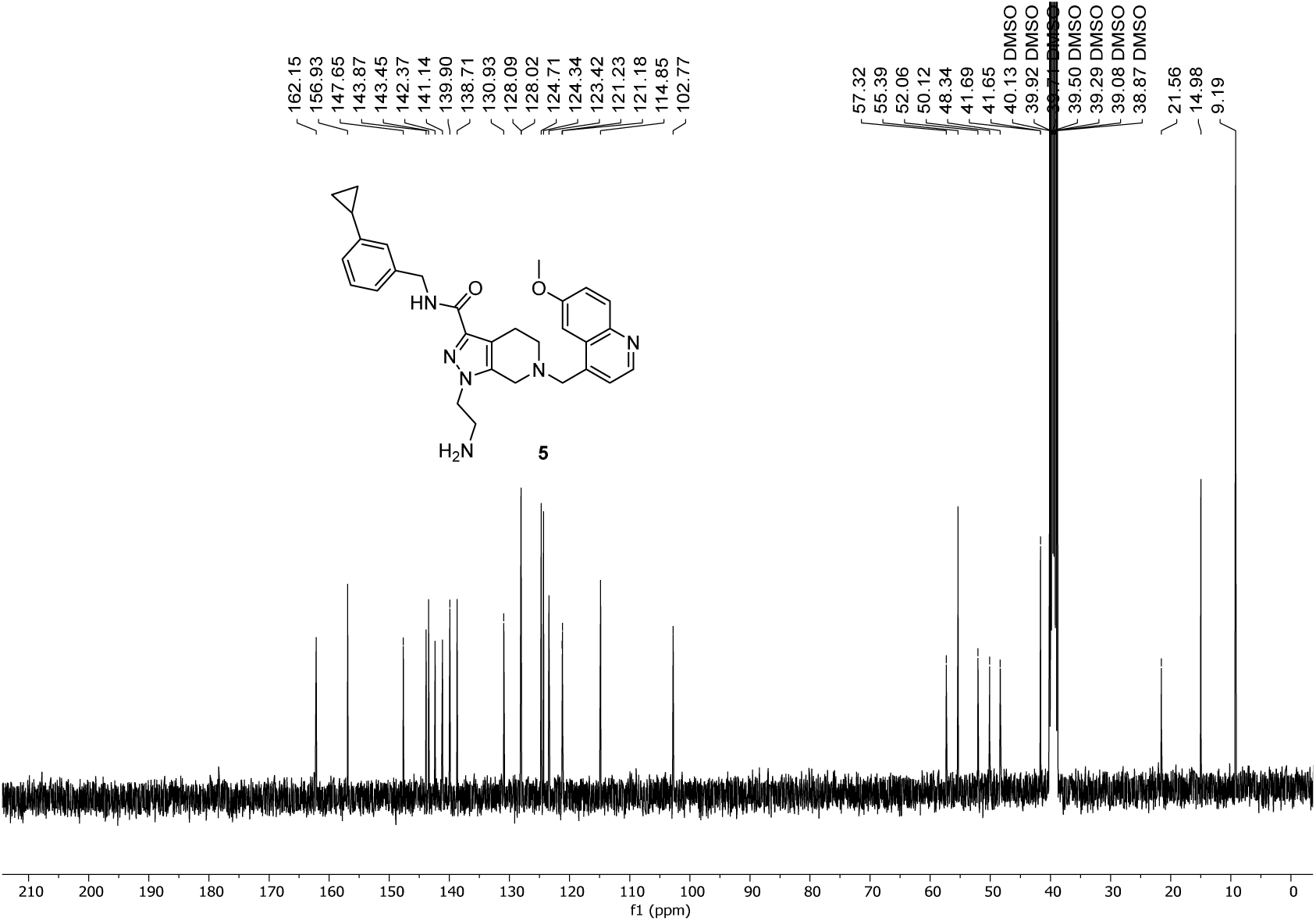

